# Assessment of Motoneuronal Regeneration and Wallerian Degeneration Following Axotomy in Postnatal Mice

**DOI:** 10.64898/2025.12.03.691768

**Authors:** Beatriu Molina-Esteve, Marina Pujol-Masip, David Ovelleiro, Jose Antonio Gomez-Sanchez, Natalia Lago, Esther Udina

**Affiliations:** Universitat Autònoma de Barcelona, Cell Biology, Physiology and Immunology, Bellaterra, Spain; Centro de Investigación Biomédica en Red sobre Enfermedades Neurodegenerativas (CIBERNED), Bellaterra, Spain; Peripheral Nervous System, Vall d’Hebron Institut de Recerca (VHIR), Vall d’Hebron Hospital Universitari, Vall d’Hebron Barcelona Hospital Campus, Barcelona, Spain; Instituto de Investigación Sanitaria y Biomédica de Alicante (ISABIAL), Alicante, Spain

**Author notes:** These authors share senior coauthorship (Natalia Lago, Esther Udina).

**Keywords:** Neonatal nerve injury, Neuronal death, Nerve regeneration, Wallerian degeneration, Transcriptomic analysis

## Abstract

Nerve injuries during early postnatal stages results in markedly different outcomes compared to adult injury, with significant motoneuronal death masking potential regenerative capacity. This study systematically evaluated motoneuronal survival, axonal regeneration, and Wallerian degeneration following peripheral nerve injury at distinct postnatal stages (P4, P10, and P30) in mice. Using ChAT-Cre/Ai9(RCL-tdT) and ChAT-Cre/RiboTag transgenic models, we assessed both histological and transcriptomic responses after sciatic nerve lesions. Injury at P4 induced substantial motoneuron death (1150%), whilst P10 and P30 animals showed minimal neuronal loss. However, when correcting for neuronal survival, P4 mice demonstrated the highest regenerative capacity, with surviving neurons achieving 100% axonal regeneration. Motoneuron-specific translatome analysis revealed that P30 animals activated a robust regeneration-associated gene (RAG) programme, including classical markers such as Atf3, Gap43, and Ngfr. In contrast, P4 neurons showed minimal RAG upregulation, suggesting they retain an intrinsic growth state that facilitates regeneration without requiring transcriptional reprogramming. P10 animals exhibited a transitional phenotype with impaired RAG activation and reduced regenerative capacity. Wallerian degeneration proceeded efficiently across all developmental stages, with age-specific differences in myelin clearance kinetics and macrophage recruitment. Transcriptomic analysis confirmed consistent downregulation of myelination programmes and upregulation of pro-regenerative markers following injury, regardless of age. These findings indicate that regenerative capacity is primarily determined by the intrinsic growth state of motoneurons rather than extrinsic factors.

**Graphical abstract:** 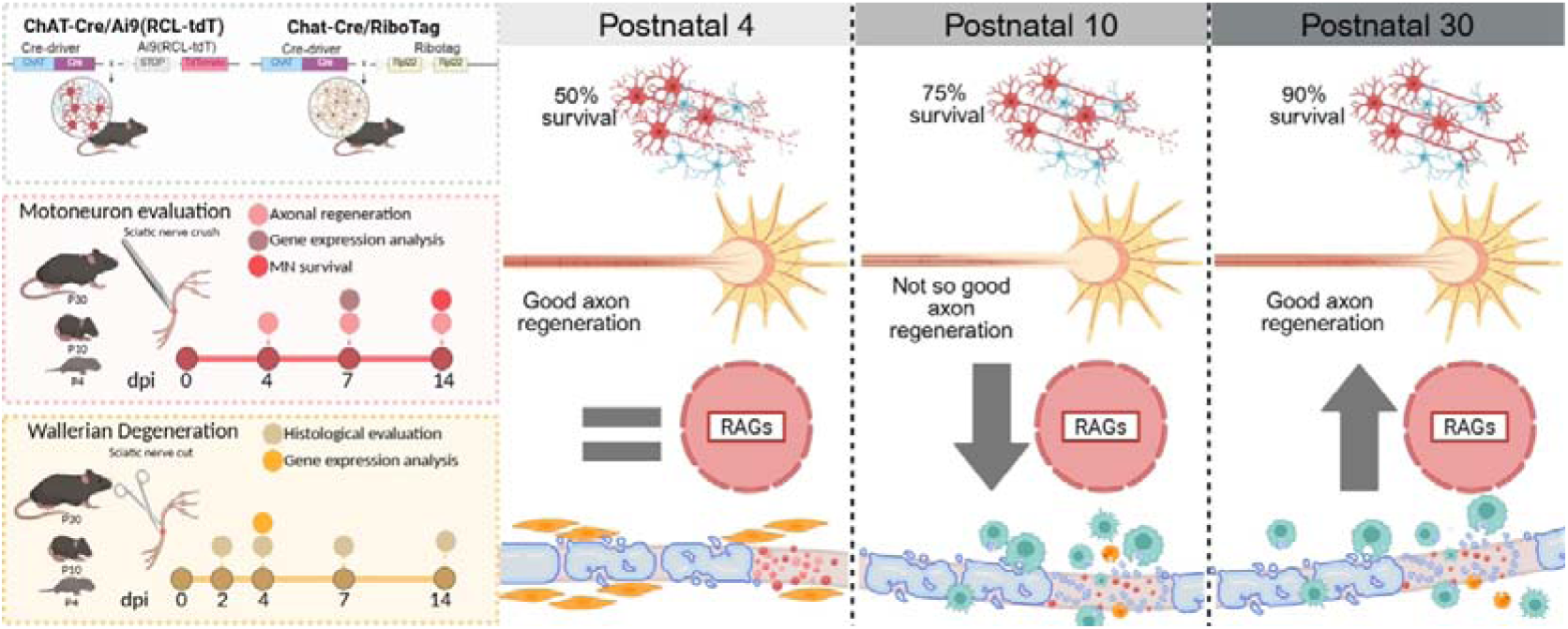

## Introduction

Lower motor neurons serve as the final effectors of the somatic motor system. Originating in the spinal cord or brainstem, their axons extend through nerves to innervate skeletal muscles. Consequently, any damage to these neurons results in muscle weakness or paralysis in the affected regions. Although the cell bodies of lower motor neurons reside within the central nervous system (CNS), they share the capacity for axonal regeneration following nerve injury with peripheral neurons. Successful peripheral nerve regeneration is attributed to two critical processes: (1) the intrinsic ability of injured neurons to switch from a neurotransmitter to a pro-regenerative state to facilitate axonal elongation and (2) the degeneration of the distal nerve stump, the so-called Wallerian degeneration, where the myelin and axonal debris at the distal stump are cleaned, and the creation of a pro-regenerative milieu provides guidance for axons to re-establish connections with their target organs (Allodi et al 2012). Activation of a genetic repair program by denervated Schwann cells (Jessen and Mirsky 2016) and resident macrophages and recruitment of hematogenous ones are key elements for successful Wallerian degeneration (Stoll and Müller 1999).

The activation of an intrinsic growth program in axotomized peripheral neurons involves the downregulation of genes related to neural activity and neurotransmission and the upregulation of several transcription factors related to growth and cytoskeleton elements (He and Jin 2016). These upregulated genes are usually known as regeneration-associated genes (RAGs). However, it is evident that neurons must survive injury to be able to switch to a pro-regenerative state. A greater susceptibility to death is observed when lesions are close to the cell body or occur at a young age. Classical works have described the marked death of motoneurons when axotomized within the first days after birth. In 1946, Romanes concluded that motoneurons are likely to die even if they are only temporarily separated from their target muscle, and this susceptibility to target deprivation diminishes with maturation, with the first six days of life being crucial for neuronal survival (Romanes 1946). Other classical studies have confirmed this massive death (Schmalbruch 1984; Li et al., 1998). A more recent paper has shown that nerve injuries in neonatal rodents induce massive loss of motoneurons but also sensory neurons (Kemp et al., 2015). Overactivation of glutamate has been implicated as a possible mechanism for motoneuron death (Iwasaki et al., 1995). The immaturity of postnatal Schwann cells has also been pointed as a contributor to this death (Komiyama and Suzuki 1992).

This massive death of neurons at postnatal stages is probably the reason why the regenerative potential of young motoneurons has been neglected in the literature. Although young neurons might have greater regenerative potential than adult neurons, this capacity can be overwhelmed by the massive neuronal loss observed at early postnatal stages.

Therefore, the aim of this study was to evaluate the regenerative capability of motoneurons after nerve injury at different postnatal stages, considering the rate of neuronal death and the genetic program activated. The degree of Wallerian degeneration of the distal stump during these different periods will also be analysed to explore the efficacy of this response at the immature stage and its impact on the regenerative process. Postnatal ages 4, 10 and 30 were selected to represent a highly vulnerable age to axotomy (P4) a transient intermediate stage between development and maturity, both for the neuron and the Schwann cells (P10) and a juvenile age which reflects onset of maturity (P30)

## Material and methods

### Mice

Postnatal 4 (P4), postnatal 10 or 11 (P10) and postnatal 30 to 35 (P30) day-old mice were used for all studies. Males and females were used in approximately equal numbers for all the groups in all the experiments. The mice were housed in a controlled environment (12-h light11dark cycle, 22 ± 2°C) in open cages with water and food ad libitum. P4 and P10 mice were housed with their mothers. The strains used in this study were acquired from Jackson Laboratory (Bar Harbor, ME, USA) and maintained in our animal facility. C57BL/6J mice were used as wild type (WT) animals. Mice expressing the fluorescent protein TdTomato in the motoneurons ChAT-Cre/Ai9 were generated by breeding homozygous Ai9 (RCL-tdT) mice (JAX stock #007909) (Madisen et al., 2010) with ChAT-IRES-Cre (choline acetyltransferase, JAX stock #006410) (Rossi et al. 2011). The same Cre-driver lines were bred to homozygous Ribotag mice (Sanz et al. 2019), resulting in mice that express hemagglutinin (HA)-tagged ribosomes in motoneurons: ChAT-Cre/Ribotag.

### Experimental design

For the evaluation of axonal regeneration and assessment of neuronal death, a sciatic nerve crush injury was applied to ChAT-Cre/Ai9(RCL-tdT) mice at P4, P10, and P30, followed by histological analysis (Figure 1a). To characterize the translatome of motoneurons after sciatic nerve crush injury at these developmental stages, RNA was isolated from injured and uninjured Chat-Cre/RiboTag mice (Figure 1b). To investigate Wallerian degeneration histology and the Schwann cell transcriptome, a sciatic nerve transection without repair was performed in WT mice at P4, P10, and P30 (Figure 1c).

**Figure 1.**
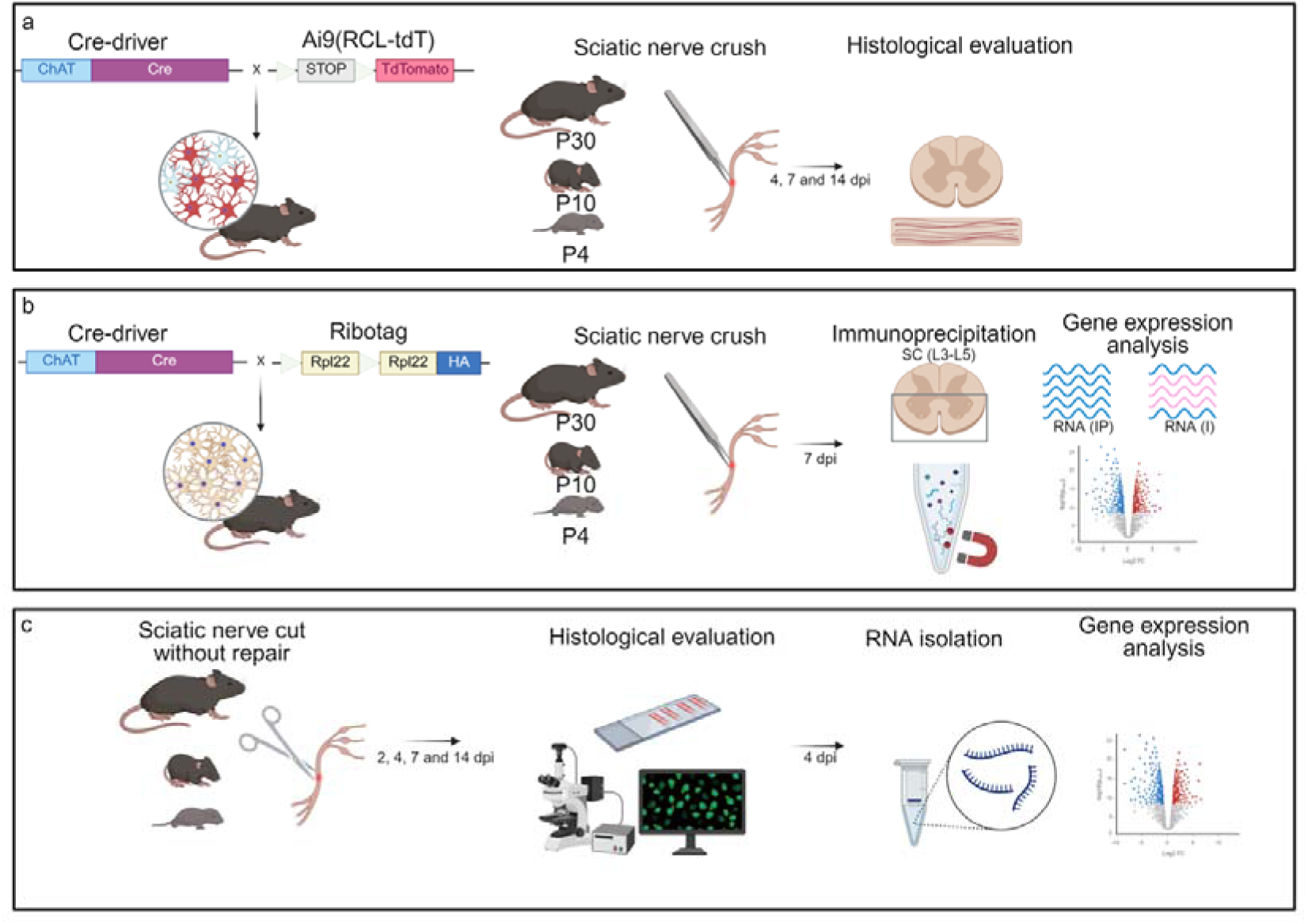
Experimental design (a) Schematic representation of the ChAT-Cre/Ai9(RCL-tdT) transgenic mice and experimental design used for the evaluation of motoneuronal survival and axonal regeneration. (b) Schematic representation of the Chat-Cre/Ribotag transgenic mice and the experimental design used for motoneuron translatome analysis. (c) Schematic representation of the experimental design used for Wallerian degeneration dynamics evaluation and sciatic nerve transcriptomics.

### Surgeries

All experimental procedures were approved by the Animal Experimentation Ethical Committee of the Universitat Autònoma de Barcelona and followed the European Communities Council Directive 2010/63/EU and Spanish National law (RD 53/2013). All surgical procedures were carried out aseptically under an operating microscope. All surgical interventions were carried out via the use of ketamine (90 mg/kg or 18 mg/kg) and xylazine (10 mg/kg or 2 mg/kg) intraperitoneally in P30 and P10 mice respectively. In P4 animals, short-term hypothermia was used to anaesthetize the pups, taking advantage of the short period needed to perform the surgery (Flecknell et al. 2015). Sciatic nerve crush was performed as follows: a mid-thigh incision was made, and the right sciatic nerve was exposed and crushed. Standard N°. Five Jeweller forceps were used for 30 seconds, resulting in axonotmesis or Sunderland grade 2 injury (Bridge et al. 1994). A 10–0 polypropylene suture was fitted into the epineurium to landmark where the crush site was at the tissue harvest. For the cut injury, the right sciatic nerve was exposed and cut proximally, resulting in complete neurotmesis or Sunderland grade 5 injury (Bridge et al. 1994). A 10–0 polypropylene suture was used to suture the proximal stump of the nerve with the muscle to prevent coaptation of the proximal and distal stumps and to avoid axonal regeneration. The skin incision was sutured with a 6–0 silk suture and disinfected with povidone iodine. All the mice were kept in a warm environment until recovery. Following surgery, all the mice were returned to their home cages and allowed to recover. During the follow-up, assessment of general health and mobility was performed daily for the first week. All the animals were sacrificed at the time of study termination under deep anaesthesia, with Dolethal (0.03 ml/30 g) administered intraperitoneally, and death was confirmed by decapitation.

### Motor axon regeneration and motoneuron death

To evaluate the number of motor axons per nerve and the number of motoneuron cell bodies 4, 7 and 14 days post injury (dpi), the ipsilateral and contralateral sciatic nerves and the lumbar spinal cord were removed, the samples were postfixed in 4% Paraformaldehyde (PFA) in phosphate buffer saline (PBS) for at least 2 hours and stored in PBS with 30% sucrose at 4°C, being post-fixed before storage. The sciatic nerves were placed on a glass slide and mounted with Fluoromount-G medium (Southern Biotech). Pressure was applied to the coverslips to flatten the nerves, and images 5 millimetres (mm) distal from the lesion point were taken via a confocal microscope (Leica SP5, 20x, z-step size of 1 μm). All intact axons were counted in stacks of 5 μm every 10 μm to avoid double counting and compared to the contralateral side. The spinal cord was serially cut (20 μm) on a cryostat (Leica), collected on glass slides, mounted with Fluoromount-G medium (Southern Biotech) and imaged with an epifluorescence microscope (Nikon Eclipse Ni, Japan) at 20x. Ipsilateral and contralateral intact motoneurons containing a clear nucleole were counted every 100 μm.

### Ribotag assay

To analyse the translatome of the injured motoneurons in the different postnatal stages, 7 days after sciatic nerve crush injury, the ventral portion of the lumbar spinal cord (L3-L5) was dissected and placed on cold Gey’s solution enriched with glucose (6 mg/mL) and homogenized in 1 mL of buffer. Age-matched uninjured animals were used as control groups (P4+7, P10+7, P30+7). After the homogenate was centrifuged, 50 μL of the supernatant was stored as an input (I) sample, whereas 5 μL of anti-HA antibody (Covance, #MMS-101R) was added to the remaining lysate and incubated for 4 hours at 4°C with rotation. Then, 200 μL of protein A/G magnetic beads (Thermo Fisher, #88803) were washed and added to the lysate overnight at 4°C with rotation. The samples were washed with high-salt buffer (HSB) to remove nonspecific binding from the immunoprecipitate (IP), and the beads were removed via a magnet. RNA was isolated from the samples via an RNeasy Micro Kit (QIAGEN, #74004) and quantified via a Quant-it RiboGreen RNA Assay Kit (Thermo Fisher, #R11490). The integrity of the RNA was assessed by using the RNA quality number (RQN), an objective metric of total RNA quality ranging from 10 (highly intact RNA) to 1 (completely degraded RNA). RQN was obtained via Lexogen CORALL library preparation - RiboCop - DA Services Bundle (Isogen Life Science B.V.). All the samples had RQN >6.

### qRT-PCR analysis

One μL of RNA was assayed via a TaqMan RNA-to-Ct 1-step Kit (Thermo Fisher, #4392938). Specific transcripts were detected via TaqMan assays: Actb (Mm02619580_g1), Fabp7 (Mm00445225_m1), and Chat (Mm01221882_m1). Relative expression was obtained by normalizing to Actb RNA levels with a standard curve method.

### Motoneuron specific RNA sequencing

Library preparation and RNA sequencing. Lexogen next-generation sequencing (NGS) services (Vienna, Austria) were used to perform RNA quality control and RNA sequencing. Libraries were prepared via the CORALL mRNA-Seq V2 preparation kit. All the samples were sequenced via the Illumina NextSeq 500 platform in runs of 2×150 bp, yielding at least 15 million reads per sample. FASTQ files were evaluated for quality via FastQC (v0.11.9). The reads were aligned to the coding DNA reference database of the mouse genome (Ensembl, GRCm39, release 103) via Salmon (v1.4.0). The quantified transcript reads were mapped to genes and imported into the R environment (v4.4.0). Genes with fewer than 100 counts across all samples were excluded from further analysis. Differential expression analysis was performed via the DESeq2 library (v1.44.0). Principal component analysis (PCA) plots were generated via the plotPCA function from DESeq2. Hierarchical clustering was performed via the pheatmap (v1.0.12) package, which focuses on the most variable genes across samples. Gene set enrichment analysis was conducted via STRINGdb and multiple databases provided by the STRING platform, including GO, KEGG, Pfam, and InterPro. Only genes with a P value<0.05 were included. Pathway analysis was performed via the Bioconductor libraries KEGGREST and pathview. The KEGG database was accessed via a RESTful web service. Pathways were analysed on the basis of genes P values via the Wilcoxon test, and graphical representations of pathways were generated, highlighting proteins by expression level.

### Histological analysis of Wallerian degeneration

To evaluate the dynamics of Wallerian degeneration in the different postnatal stages 2, 4, 7 or 14 dpi, the ipsilateral and contralateral sciatic nerves were removed, fixed in 4% PFA in PBS for at least 2 hours and stored in PBS supplemented with 30% sucrose at 4°C. Longitudinal sciatic nerve sections of 8-µm thickness obtained with a cryostat (Leica, Wetzlar, Hermany) were subjected to Luxol fast blue staining (LFB, Sigma; St. Louis, MO, USA) for myelin clearance quantification. Following gradual dehydration, the sections were immersed in a 1 mg/mL LFB solution in 95% ethanol and 10% acetic acid overnight at 37°C. The sections were subsequently rinsed in distilled water, destained in a solution of 0.05% LiCO3 in distilled water for 10 seconds and finally rinsed in 70% ethanol. The sections were then dehydrated and mounted with DPX mounting medium (Sigma; St. Louis, MO, USA). Images were acquired at 20x via an epifluorescence microscope under bright light (Nikon Eclipse Ni, Japan). For lipid accumulation quantification, the sections were incubated in Oil Red O (ORO) solution for 10 minutes at room temperature and then placed under running tap water for 30 minutes. The sections were then mounted with Fluoromount-G medium (Southern Biotech). Images were acquired at 20x via an epifluorescence microscope under bright light (Nikon Eclipse Ni, Japan). For phagocyting macrophage density quantification, samples were permeabilized with PBS and 0.3% Triton X-100 (PBST) and blocked with 1.5% normal donkey serum (NDS) (Vector Laboratories). The sciatic nerve slices were incubated overnight at 4°C with a rat anti-CD68 antibody (1:500, Bio-Rad). The slices were washed and incubated with secondary anti-rat antibody (Alexa Fluor 488, Invitrogen) for 2 h at room temperature. After washing, the slides were mounted with Fluoromount-G medium (Southern Biotech). Images were acquired at 20x via an epifluorescence microscope (Nikon Eclipse Ni, Japan). To analyse the percentage of degenerating fibres in the distal nerve after a sciatic nerve cut without repair, semithin sections of the distal nerves were obtained at 2, 4 and 7 dpi and postfixed in glutaraldehyde11paraformaldehyde (3%/3%) in cacodylate buffer solution (0.1 M, pH 7.4). The samples were postfixed in 2% Osmium Tetroxide, dehydrated through ethanol series, and embedded in epoxy resin embedding medium (Epon) resin. Transverse semithin sections (0.5 μm) were stained with toluidine blue and examined via light microscopy.

To quantify myelin loss over time in the different postnatal stages postinjury, the percentage of the area stained with LFB, ORO and anti-CD68 immunostaining was measured via Fiji/ImageJ software.

### Sciatic nerve RNA analyses

To analyse the transcriptomics of the injured sciatic nerves in the different postnatal stages 4 days after cut injury, the ipsilateral and contralateral sciatic nerves were extracted and snap frozen until RNA extraction. RNA from the sciatic nerve was isolated via the RNeasy Micro Kit (QIAGEN, #74004). The contralateral nerves served as the control group.

### Library preparation and RNA sequencing

BGI Genomics Co., Ltd. (Shenzhen, China) performed the RNA quality control and RNA sequencing. Libraries were prepared via the DNBSEQ Eukaryotic Strand-Specific mRNA Library Kit. All samples were sequenced via the DNBSEQ (DNBSEQ Technology) platform in runs of 2x150 bp, yielding at least 20 million reads per sample. FASTQ files were evaluated for quality via FastQC (v 0.11.9). The reads were aligned to the coding DNA reference database of the mouse genome (Ensembl Mouse database, Genome assembly: GRCm39.cdna.all, release 113) via Salmon (v1.10.3). The “DESeq2” library (v 1.48.0) was used to perform the differential expression analysis pipeline. The quantified transcript reads were mapped to genes and imported into the R environment (R version 4.5.0). Only genes with more than 10 counts in the smallest group were used in the differential analysis. PCA and uniform manifold approximation and projection (UMAP) plots were generated via the plotPCA function from DESeq2. Hierarchical clustering was performed via the pheatmap (v1.0.12) package, which focuses on the 80 most variable genes across samples. Gene set enrichment was performed via the R library “STRINGdb” 7 via the String 8 platform, which includes GO, PubMed, STRING, KEGG and Reactome, among others. Only proteins with a P value<0.05 were included. Pathway analysis was performed for each comparison. The bioconductor libraries “KEGGREST” (v 1.48) and “pathview” (v 1.48) were used. The KEGG database is accessed in this process via a RESTful web service via the R package. Pathways were analysed on the basis of gene P values via the Wilcoxon test, and graphical representations of pathways were generated, highlighting genes by expression level. In addition to the differential expression obtained, a single time series involving both the “Ctrl” and “Inj” samples at the three times (4, 10 and 30) has been modelled by designing a formula that models the type of sample and the first time point, differences over time, and any type-specific differences over time, followed by a likelihood ratio test and the removal of type-specific differences over time.

### Data analysis

GraphPad Prism 9 (version 9.0.1) was used for statistical analysis. The normal distribution of the samples was confirmed with the Shapiro11Wilk test (p>0.05). The percentages of regeneration and neuronal death were analysed via two-way ANOVA and paired t tests, respectively, followed by the Sidak’s test. Histological statistical analyses were performed via two-way ANOVA followed by the Sidak’s test. The total number of axons was analysed via the Kruskal11Wallis’s test followed by Dunn’s test. The percentage of myelinated/degenerating axons was analysed via two-way ANOVA followed by the Sidak’s test. Differences were considered statistically significant if p<0.05. All the data are expressed as the group mean ± standard error of the mean (SEM).

## Results

### Response of motoneurons to nerve injury at different postnatal stages

#### Evaluation of neuronal death and axonal regeneration

Motoneuron death was evaluated 14 dpi by counting the total number of neuronal cell bodies present in the ipsilateral ventral horn of the lumbar portion of the spinal cord (L3-L5) and comparing them to those on the contralateral side (Figure 2a). On the contralateral side, approximately 12 motoneurons/slice were found at all the ages evaluated. When analysing the ipsilateral side, we observed a significant reduction when animals were injured at P4 (6 motoneurons/slice) and a non-significant reduction at P10 and P30 (9 and 8 motoneurons/slice respectively; Figure 2b). Whereas injuries at P10 and P30 resulted in 75% and 80% of motoneuron survival at 14 dpi, injuries at P4 resulted in only 50% survival. This reduced survival when injury is applied at P4 mice is significant compared to P10 (p<0.05) and P30 mice (p<0.005). (Figure 2c)

**Figure 2.**
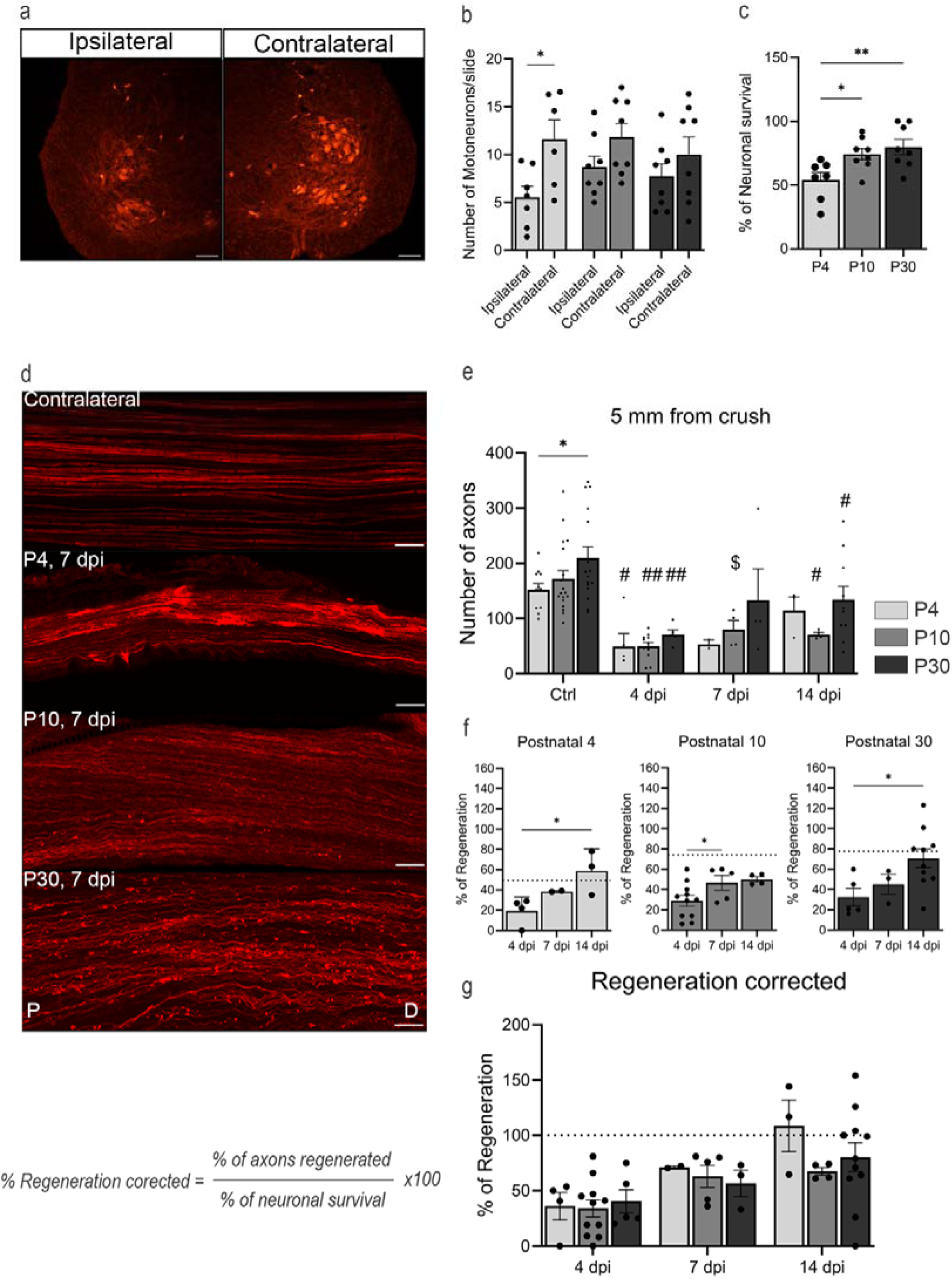
(next page). Histological characterization of the motor neuron response after peripheral nerve injury at different postnatal stages. (a) Representative images of the ipsilateral and contralateral ventral horns of the lumbar portion of the spinal cord of P4 mice at 14 dpi, showing the ChAT+ motoneurons in red. Scale bar=100 µm. (b) Quantification of the average number of motoneurons in the lumbar spinal cord on both the ipsilateral and contralateral sides at 14 dpi across different postnatal stages. Statistical analysis was performed via two-way ANOVA followed by Sidak’s correction test. *p<0.05 (c) Percentage of surviving motoneurons at 14 dpi in the different postnatal stages. Statistical analysis was performed via one-way ANOVA followed by Tukey’s correction test. *p<0.05; **p<0.001. (d) Representative longitudinal sections of the distal portion of the sciatic nerve at 7 dpi in P4, P10, and P30 mice. Scale bar: 50 µm. P: proximal; D: distal. (e) Total number of axons 5 mm distal to the lesion site in P4, P10, and P30 mice in both control and crushed sciatic nerves at 4, 7, and 14 dpi. Statistical analysis was performed via two-way ANOVA followed by Sidak’s correction test. *p<0.5; $ p<0.5 vs. control; p<0.01 vs. control; ##. p<0.0001 vs. control. (f) Percentage of regenerated axons at 4, 7, and 14 dpi in P4, P10, and P30 mice relative to the contralateral side. The dotted line represents the percentage of surviving motoneurons at 14 dpi. Statistical analysis was performed via one-way ANOVA followed by Tukey’s correction test. *p<0.05 (g) Regeneration index corrected for neuronal survival at 4, 7, and 14 dpi in P4, P10, and P30 mice relative to the contralateral side. Statistical analysis was performed via two-way ANOVA followed by Sidak’s correction test.

When we analysed the number of motor axons in intact nerves at different postnatal stages (Figure 2d, e), we observed a significantly greater number of axons in P30 mice than in P4 mice (p<0.05). After crush injury at different postnatal stages, we evaluated the number of motor axons growing 5 mm distal to the injury site at 4, 7 and 14 dpi. As expected, in all the experimental groups, we observed a significant reduction in the number of axons 5 mm distal to the injury at 4 dpi compared with those on the contralateral side (p<0.05). Interestingly, at 14 dpi, only P4 mice reached control values (p<0.05), indicating good axonal regeneration at this stage. However, since the basal (control) number of axons differed between postnatal stages, we normalized the number of regenerating axons distal to the lesion versus the number of intact axons in the control nerves to obtain the percentage of regenerated axons (Figure 2f). Considering this correction, when injury was applied at P4 and P10, we observed that approximately 55% of the axons regenerated at 14 dpi, a percentage that increased to 75% when injury was applied to P30 mice. However, since the death of motoneurons directly impacts the number of axons that may regenerate after injury, we also applied a correction factor to normalize the regeneration data considering the survival of motoneurons at different ages. This correction factor was calculated as follows:

Percentage of regenerated axons/percentage of motoneuron survival *100.

When this correction factor was applied (Figure 2g), only P4 mice reached 100% of regeneration. Although there were no significant differences between the groups in terms of these corrected regeneration rates, motoneuron regeneration in P4 animals was considerably good, despite the high rate of neuronal loss observed.

### RNA isolation from ChAT-Cre/Ribotag is motoneuron specific

To further study whether the response of motoneurons to injury differed among the different postnatal stages, we used the Ribotag assay to isolate translated mRNA from motoneurons at different postnatal ages. By breeding ChAT-Cre driver mice with Ribotag mice, we could target motoneuron ribosomes, which were tagged with HA, as previously validated in our laboratory (Bolívar et al. 2024). By harvesting the ventral portion of the lumbar spinal cord, we could specifically isolate the ribosomes of the motoneurons. Following RNA isolation, RT11qPCR analysis of the IPs revealed marked enrichment of the motoneuron-specific transcript Chat (choline acetyltransferase), whereas the glial transcript Fabp7 (fatty acid binding protein 7) was notably depleted across all IPs (Figure 3a). This result confirms the specificity of ribosome isolation to neuronal populations within our samples.

**Figure 3.**
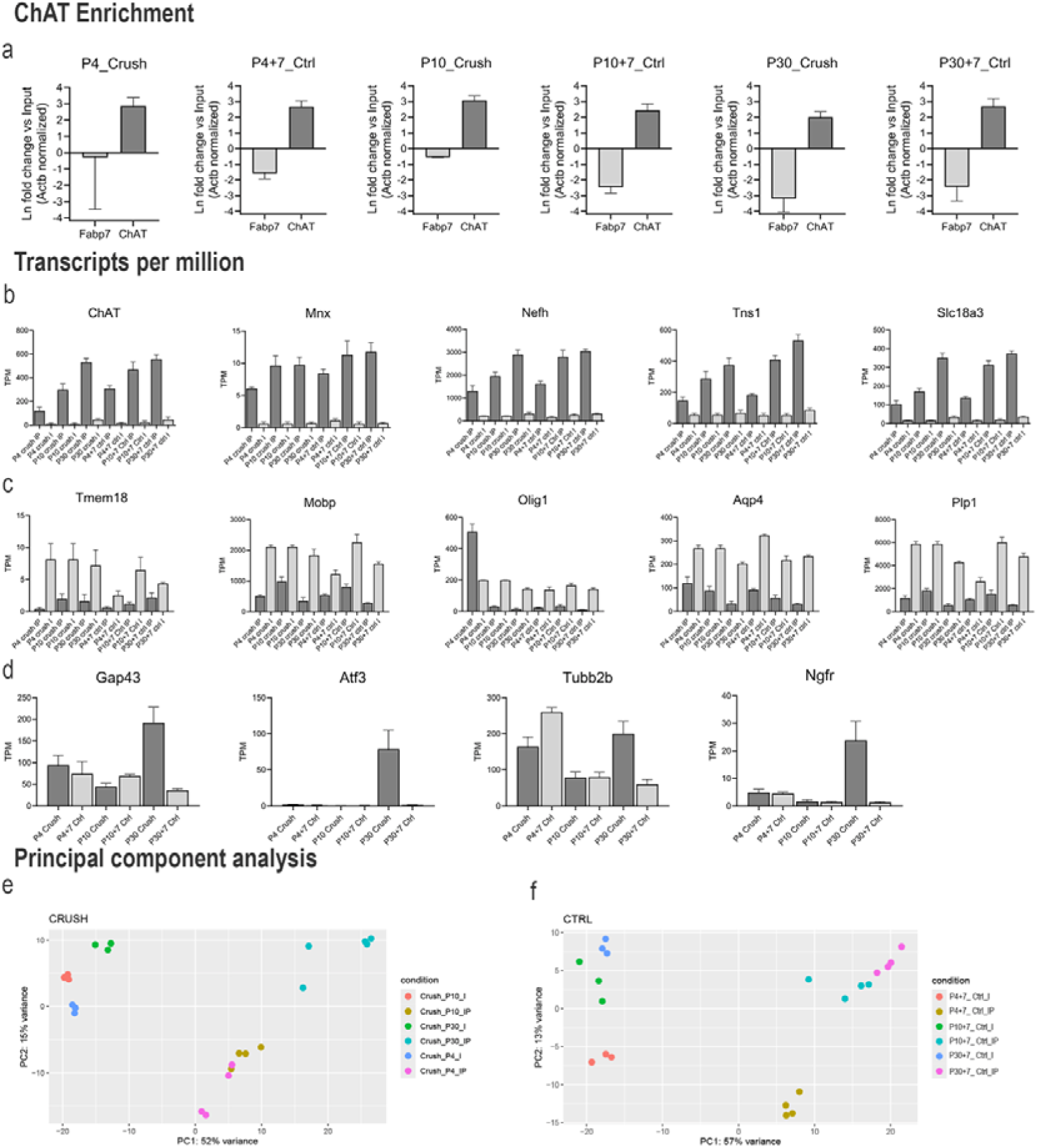
Validation of the specificity of the ChAT-Cre/Ribotag mice. (a) RT12qPCR revealed enrichment of cell type-specific transcripts in the IPs of ChAT and depletion of the glial transcript Fabp7. (b) TPM of motoneuron-specific genes expressed in the IP and I. (c) TPM of glial-specific genes in both the IP and I. (d) TPM of regeneration-specific genes expressed in the crush and control samples. (e, f) PCA of the crush and control samples included in the study.

RNA sequencing revealed differential expression of over 15,000 genes across each group of interest. To validate the dataset, we examined the transcripts per million (TPM) of several cell type-specific markers. Motoneuron IPs were enriched for established motoneuron markers, including ChAT, Mnx, Nefh, Tns1, and Slc18a3 (Figure 3b). In contrast, transcripts associated with glial genes, such as Tmem119, Mobp, Olig1, Aqp4 and Plp1, were depleted in the IPs, confirming their cell type specificity (Figure 3c). RAGs, including Gap43, Atf3, Tubb2b, and Ngfr, were enriched in most of the injured samples compared with the control samples (Figure 3d). Principal component analysis further demonstrated clear clustering of samples on the basis of their experimental group, supporting the robustness of the dataset (Figure 3e, f).

### Pathway enrichment analysis

To investigate the molecular response of motoneurons to sciatic nerve crush at different postnatal stages, we performed pathway enrichment analysis (Figure 4).

**Figure 4.**
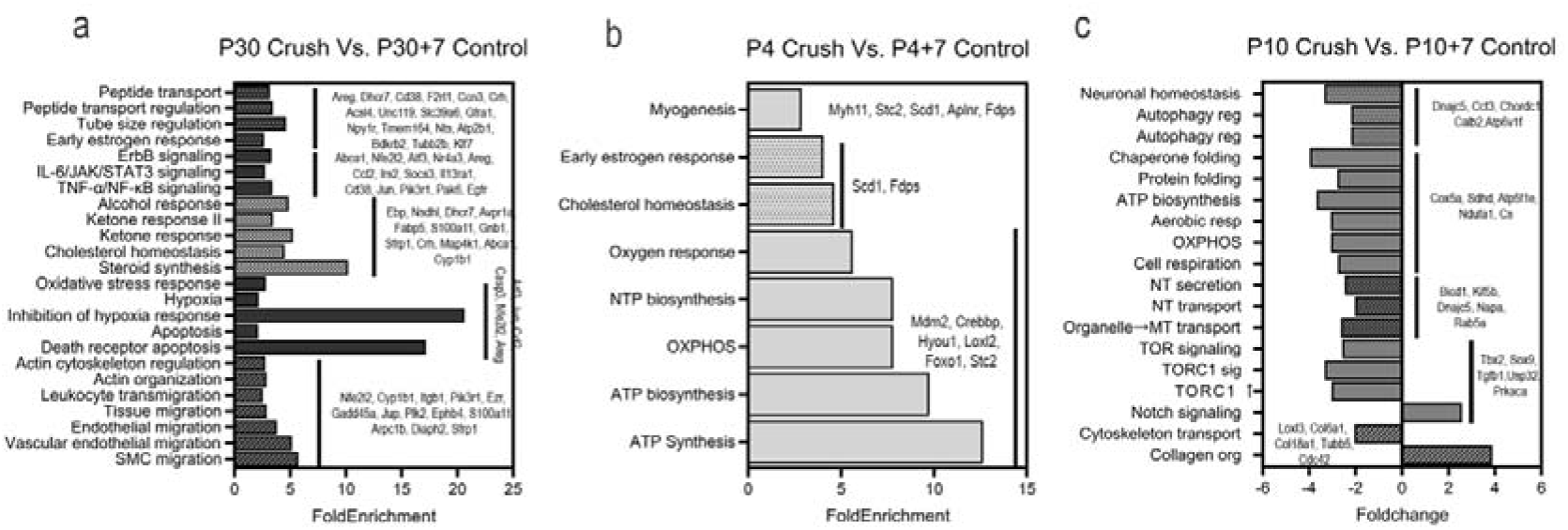
Pathway enrichment analysis. (a) Molecular pathways enriched in P30 crush animals compared with P30+7 control animals. (b) Molecular pathways enriched in P4 crush animals compared with P4+7 control animals. (c) Molecular pathways enriched and downregulated in P10 crush animals compared with those in P10+7 control animals.

The analysis revealed distinct stage-dependent profiles. In P30 mice (Figure 4a), we selected 24 enriched pathways; most of them are related to migration and cellular organization, stress response, lipidic metabolism and inflammatory response. In P4 injured mice compared to uninjured P4+7 mice (Figure 4b), the most enriched pathways are related to mitochondrial energetic metabolism and early regulatory processes. In injured P10 mice compared to uninjured Postnatal 10+7 mice (Figure 4c), most of the pathways are downregulated, including those related to neuronal homeostasis, autophagy, energetic metabolism, and cytoskeleton reorganization.

### Gene Set Enrichment Analysis (GSEA)

To investigate whether sets of genes exhibited a consistent increase or decrease in expression across different postnatal stages following injury, we performed gene set enrichment analysis (GSEA). Comparisons were made between the injured group and the corresponding age-matched control group (Injured P4 vs. Intact P4+7, Injured P10 vs. Intact P10+7, Injured P30 vs. Intact P30+7). Using our custom list of RAGs and RStudio software (RStudio Team, 2024, version 2024.12.0+467), we conducted GSEA (Figure 5a-c). We subsequently focused on the leading-edge RAGs identified via GSEA to pinpoint the genes most significantly enriched at each postnatal stage. In P30 animals, several classical RAGs, including Atf3, Ccl2, Ngfr, Gap43, and Adcyap1, were upregulated compared with their noninjured counterparts. In contrast, compared with noninjured animals of the same age, injured P4 animals did not exhibit any significantly upregulated RAGs. Interestingly, injured P10 mice presented a downregulation of several classical RAGs, such as Rac1 and Stat3, relative to noninjured P10+7 animals. Following the GSEA findings, we next examined the expression levels of the identified leading-edge RAGs by assessing their TPM values across the experimental groups (Figure 5 d-i). Consistent with the GSEA results, injured P30 animals presented visibly greater expression of these RAGs than did their intact P30+7 counterparts. In contrast, the TPM values for these RAGs in both injured P10 and injured P4 animals showed minimal discernible changes relative to their respective age-matched intact control groups.

**Figure 5.**
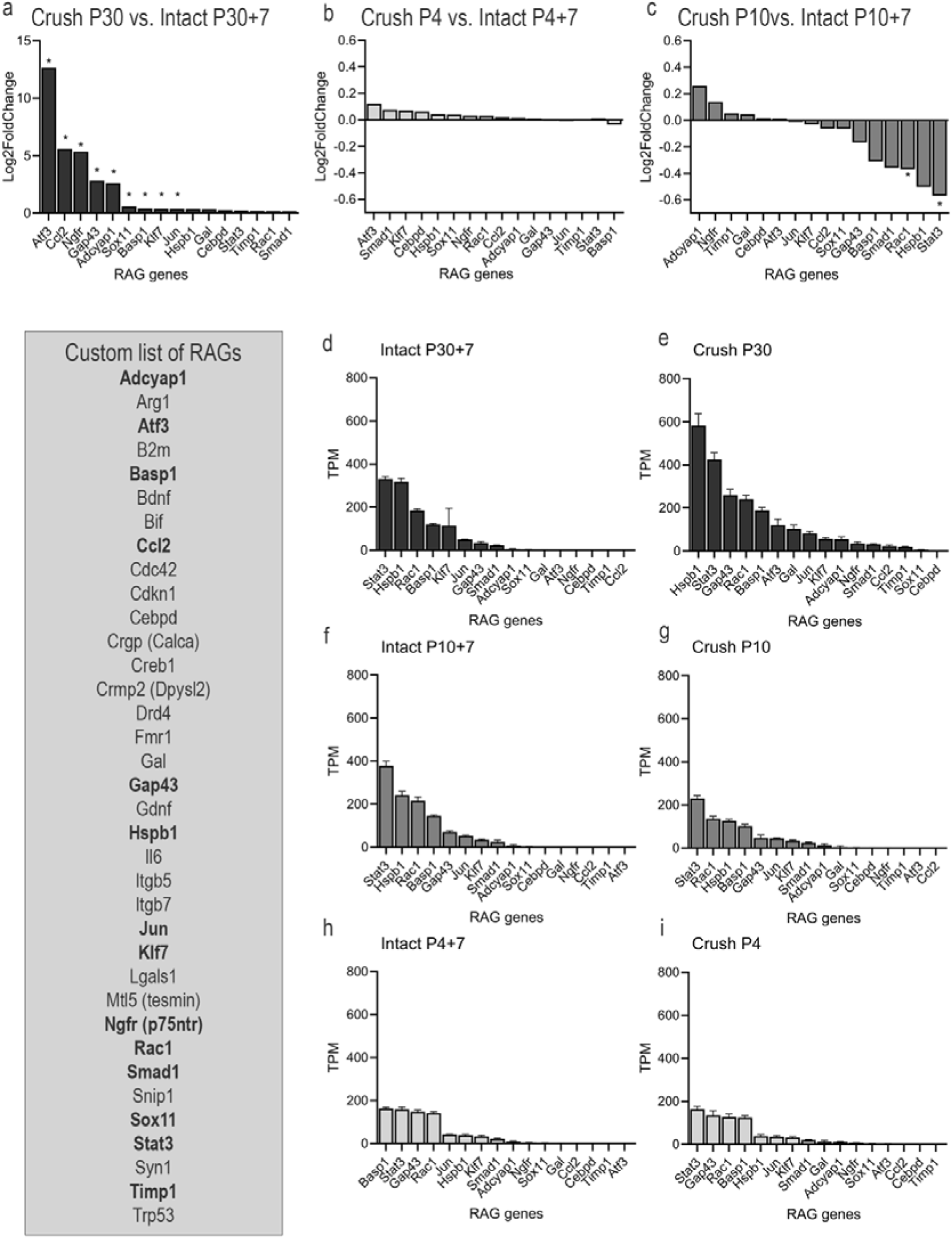
(next page). Expression of RAGs at different postnatal stages. (a) Log₂Fold change in the number of enriched RAGs in motor neurons from injured P30 mice compared with uninjured P30+7 controls. (b) Log₂Fold change in the number of enriched RAGs in motor neurons from injured P4 mice, analysed at P11, compared with that in age-matched uninjured P4+7 controls. (c) Log₂Fold change in the number of enriched RAG genes in motor neurons from injured P10 mice, analysed at P17, compared with those in uninjured P10+7 controls. Differential expression and enrichment analyses in (a–c) were performed via gene set enrichment analysis (GSEA) in R. Gene sets were considered significantly enriched at false discovery rate (FDR)<0.05. (d–i) Transcripts per million (TPM) of selected RAGs in motor neurons injured at P30, P10 and P4 mice and age-matched controls. The data are presented as the means ± SEMs (n=4 mice per group).

### Characterization of Wallerian Degeneration at Different Postnatal Stages

To evaluate Wallerian degeneration, a cut injury was applied to the sciatic nerve at different postnatal stages. This was performed to isolate the distal segment from the proximal segment, thereby preventing axonal regeneration into the distal segment.

### Histological characterization of Wallerian degeneration

LFB staining (Figure 6a) was used to evaluate myelin integrity in the sciatic nerve following peripheral nerve injury (PNI) at different postnatal stages. In contralateral nerves from injured animals, LFB staining consistently revealed a homogeneous and intense blue signal, indicative of intact and well-organized myelin sheaths. In contrast, ipsilateral nerves from animals at all ages examined (P4, P10, and P30) presented altered LFB staining patterns characterized by reduced staining intensity and increased heterogeneity, suggesting progressive demyelination post injury. Quantification of the LFB-labelled area (Figure 6b) revealed robust myelin content in the contralateral nerves of P30 mice, which was significantly lower in P4 and P10 mice. Following injury, myelin clearance in P30 mice gradually occurred, with maximal loss observed at 7 dpi. In P10 mice, myelin clearance also progressed over time, reaching a peak at 4 dpi, suggesting a more rapid response than at P30. Notably, P4 mice exhibited an absence of detectable myelin as early as 2 dpi, with levels remaining consistently low through 14 dpi. These results indicate that while P10 and P30 mice follow a similar profile of progressive myelin clearance, P4 mice display an atypical response, likely reflecting developmental immaturity and distinct cellular mechanisms engaged following injury at this early postnatal stage. ORO staining (Figure 7a) was employed to assess lipid droplet accumulation resulting from myelin degradation following PNI at different postnatal stages. In sciatic nerve tissue from uninjured control animals, a uniform and faint reddish staining pattern was consistently observed. In contrast, injured nerves from all age groups exhibited a heterogeneous staining pattern characterized by the presence of discrete, intensely red lipid aggregates along the nerve fibres and within the surrounding connective tissue. Quantification of the ORO-labelled area (Figure 7b) in contralateral nerves revealed no lipid accumulation at any postnatal stage. Following injury, P30 mice presented a progressive increase in lipid accumulation, peaking at 7 dpi. In P4 mice, lipid droplets were observed at 4 dpi but rapidly decreased by 7 and 14 dpi. P10 mice displayed a gradual accumulation of lipids, reaching a peak at 4 dpi, followed by a progressive decline through 14 dpi. These findings suggest that lipid accumulation dynamics following PNI vary with developmental stage, with P10 and P30 mice exhibiting more sustained lipid accumulation than transient accumulation, as observed in P4 mice.

**Figure 6.**
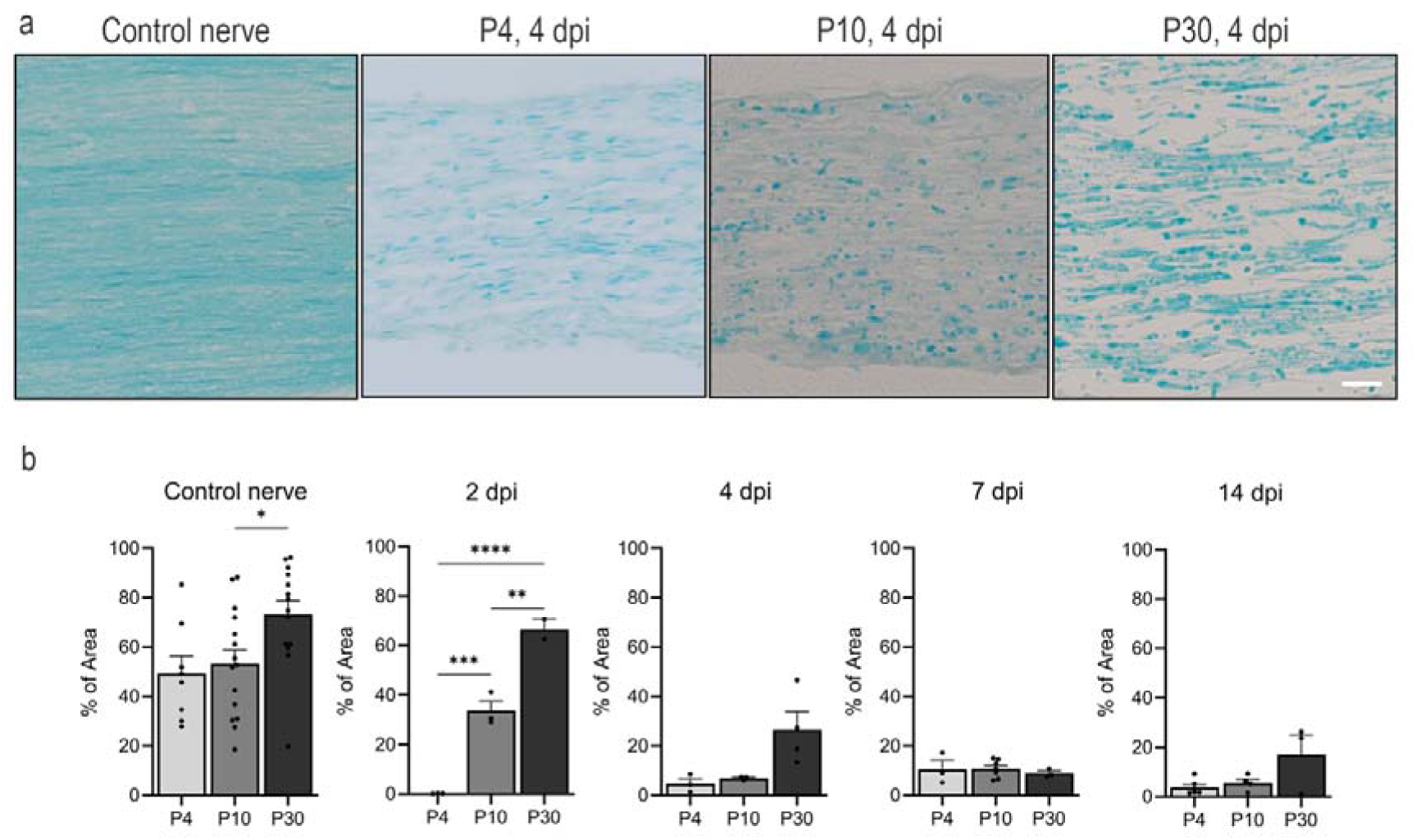
Histological evaluation of myelin clearance dynamics in different postnatal stages. (a) Representative images of LFB staining of the distal portion of the sciatic nerve at different postnatal stages after a cut lesion at 4 dpi and contralateral to it. (b) Percentage of the area stained with myelin in the distal portion of the different postnatal sciatic nerves (P4, P10 and P30) after a cut injury without repair at 2, 4, 7 and 14 dpi. (n= 3127 animals per group) * p<0.05, **p<0.01, *** p<0.001, **** p<0.0001 calculated by two-way ANOVA followed by Bonferroni correction for multiple comparisons. Scale bar: 100 µm.

**Figure 7.**
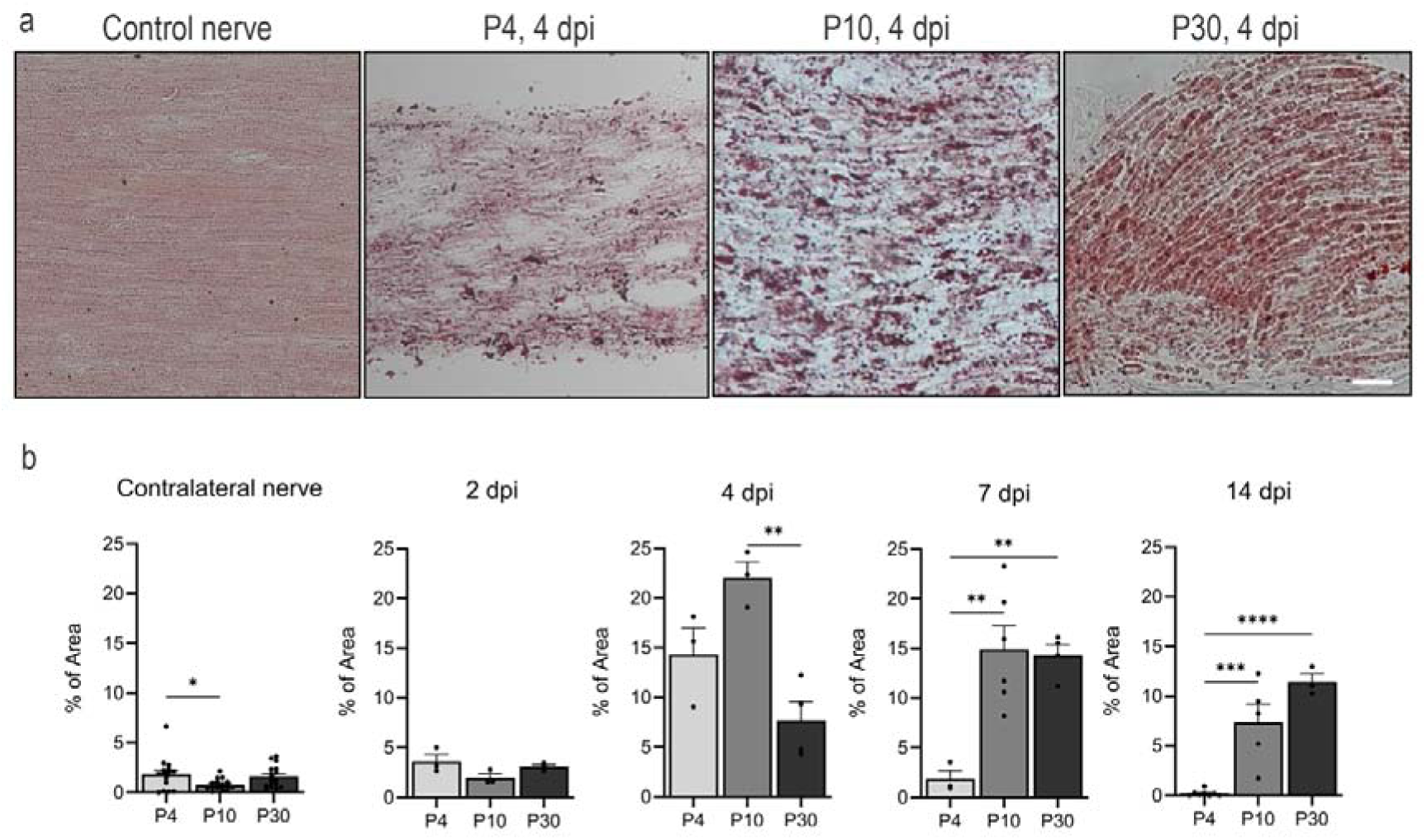
Histological evaluation of lipid accumulation in different postnatal stages. (a) Representative images of ORO staining of the distal portion of the sciatic nerve at different postnatal stages after a cut injury at 4 dpi and contralateral to it. (b) Percentage of the area stained with lipid droplets in the distal portion of the different postnatal sciatic nerves (P4, P10 and P30) after a cut injury without repair at 2, 4, 7 and 14 dpi. (n= 3127 animals per group) *p<0.05, **p<0.01, *** p<0.001, **** p<0.0001 calculated by two-way ANOVA followed by Bonferroni’s correction for multiple comparisons. Scale bar: 100 µm.

To quantify the degree of macrophage recruitment in the sciatic nerve following PNI across different postnatal stages, immunofluorescence staining for CD68 was performed (Figure 8a). In contralateral nerves from injured animals, only sparse, small CD68-positive cells—consistent with resident macrophages—were observed. In contrast, injured nerves from P10 and P30 mice presented a marked increase in macrophage density, with cells exhibiting a larger and rounder morphology characteristic of phagocytic activity. Quantification of the CD68-labelled area (Figure 8b) in contralateral nerves confirmed the absence of macrophage recruitment and active phagocytosis across all postnatal stages. After injury, macrophage recruitment in P30 mice increased progressively over time, peaking at 14 dpi. P10 mice displayed a similar trend, with maximal macrophage accumulation at 7 dpi followed by a reduction at 14 dpi. Notably, P4 mice did not exhibit a significant increase in the number of macrophages at any time point. These results indicate that macrophage recruitment and activation following PNI are developmentally regulated, with P4 nerves showing an absence of macrophage recruitment, whereas P10 and P30 nerves mount more robust and temporally distinct macrophage responses.

**Figure 8.**
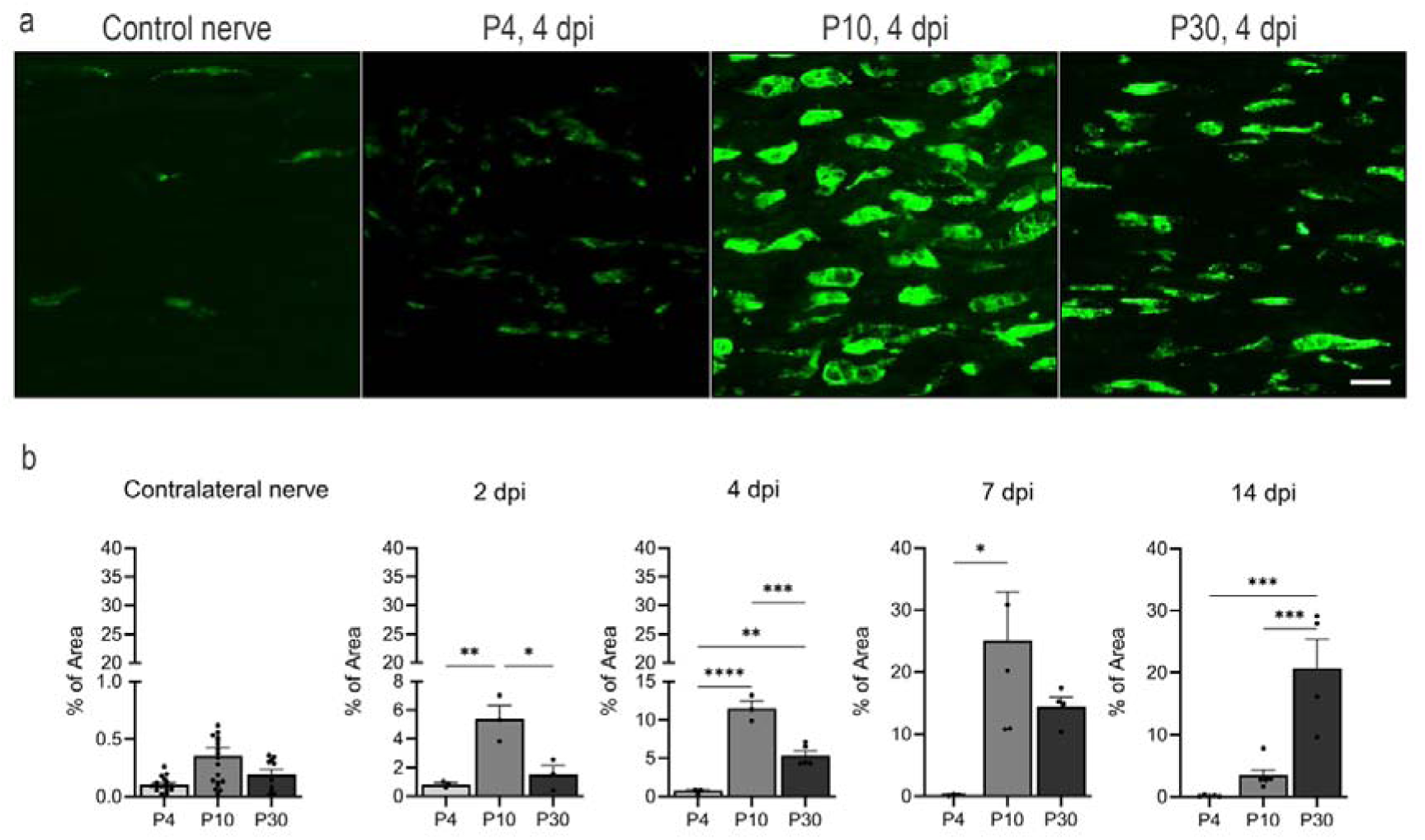
Histological evaluation of the activity of phagocytosing macrophages in different postnatal stages. (a) Representative images of anti-CD68 immunostaining in the distal portion of the sciatic nerve at different postnatal stages after a cut injury at 4 dpi and contralateral to it. (b) Percentage of the area stained with phagocytosing macrophages in the distal portion of the different postnatal sciatic nerves (P4, P10 and P30) after a cut injury without repair at 2, 4, 7 and 14 dpi. (n= 3127 animals per group) *p<0.05, **p<0.01, ***p<0.001, ****p<0.0001 calculated by two-way ANOVA followed by Bonferroni’s correction for multiple comparisons. Scale bar: 100 µm.

Structural analysis of transverse sciatic nerve sections (Figure 9a) revealed the presence of myelinated axons in the contralateral nerves across all postnatal stages. In uninjured nerves (Figure 9b), myelinated axons were readily identifiable by their characteristic circular to oval morphology, with dense, compact myelin sheaths surrounding lighter central axoplasmic regions. The myelin appeared uniform in both thickness and organization, which was consistent with intact and mature nerve fibres. In nerves from uninjured P4 mice, we also observed axons encased by Schwann cells with a diameter compatible with myelinated fibres but lacking a clearly defined myelin sheath. These structures likely represent axons undergoing early stages of myelination (Supplementary figure 1). Quantification of the total number of myelinated axons (Figure 9c) revealed an increase in axon number with increasing postnatal age. The injured (ipsilateral) nerves of P10 and P30 mice (Figure 9d) revealed a gradual reduction in the number of myelinated axons over time. In contrast, injured P4 nerves exhibited a complete absence of myelinated axons from 2 to 7 dpi. Injured nerves displayed hallmark features of Wallerian degeneration (Figure 9e), including disorganized tissue architecture, myelin sheath fragmentation, and axonal swelling. The myelin rings appeared collapsed, folded, or partially degraded, and numerous vacuolated axonal profiles and myelin debris were scattered throughout the tissue (white arrows), which was consistent with ongoing myelin breakdown and axonal degeneration. The quantification of degenerating axons in the ipsilateral nerves (Figure 9f) revealed a consistent temporal pattern across all postnatal stages: the number of degenerating axons peaked at 4 dpi. This peak was less clear in P30 animals, with similar numbers of degenerating axons at 4 and 7 dpi.

**Figure 9.**
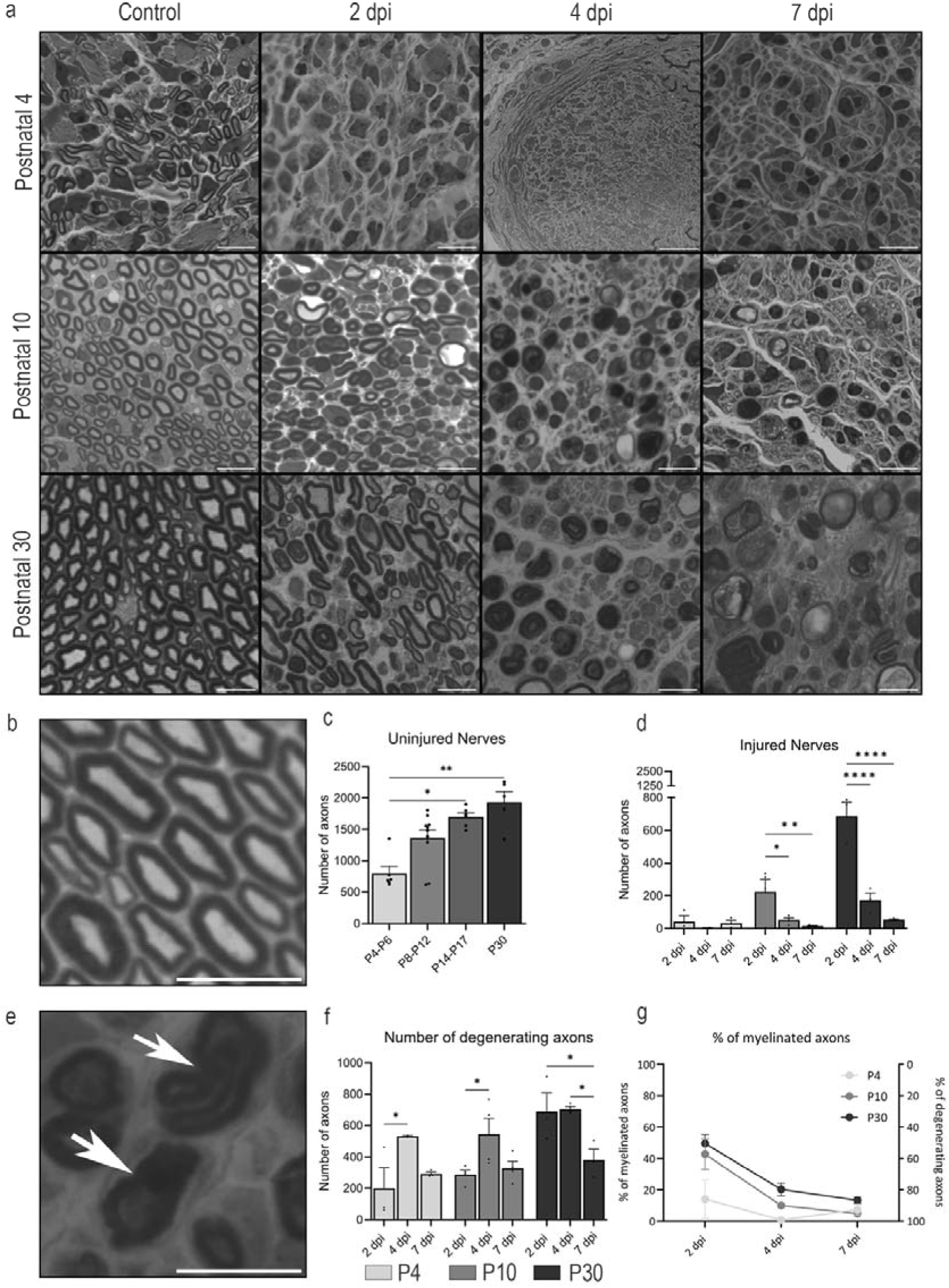
(next page). Ultrastructural analysis and axonal density dynamics following sciatic nerve injury across postnatal stages. (a) Representative images of transverse sciatic nerve sections from uninjured (control) and injured (2, 4, and 7 dpi) P4, P10, and P30 mice. (b) Representative image of axons surrounded by myelin sheaths. (c) Quantification of the total number of myelinated axons in control sciatic nerves at different postnatal stages. (d) Quantification of the total number of myelinated axons in injured sciatic nerves at different postnatal stages. (e) Representative image of different degenerating axons after PNI. (e) Quantification of the total number of degenerating axons in injured sciatic nerves at different postnatal stages. (e) Percentages of myelinated and degenerating axons in the different postnatal stages over time. Statistical significance was determined by the Kruskal12Wallis’s test or two-way ANOVA followed by Tukey’s correction for multiple comparisons; * p<0.05, ** p<0.01 ****p<0.0001. Scale bar: 10 µm

The number of degenerating axons with intact nerves was normalized to calculate the percentage of degenerating axons found over time in the different postnatal stages (Figure 9g). In P10 and P30 mice, ∼50% of the axons were degenerating at 2 dpi, with degeneration peaking at 7 dpi. Interestingly, in P4 mice, ∼85% of the axons had already degenerated at 2 dpi, reaching a peak of degeneration at 4 dpi, suggesting a more rapid degenerative response in the most immature nerves.

Transcriptomic analyses from injured sciatic nerves during the postnatal stages To investigate the transcriptomic differences underlying Wallerian degeneration at different postnatal stages, we performed RNA-seq on sciatic nerves from injured P4, P10, and P30 mice. Age-matched uninjured nerves served as controls. The dataset included 5 samples per group, with sequencing yielding an average of ∼23.5 million paired end reads per sample. PCA analysis (Figure 10) shows clustering of the samples based on their gene expression profiles. The left and right panels display the control (Ctrl) and injured (Inj) samples, respectively.

**Figure 10.**
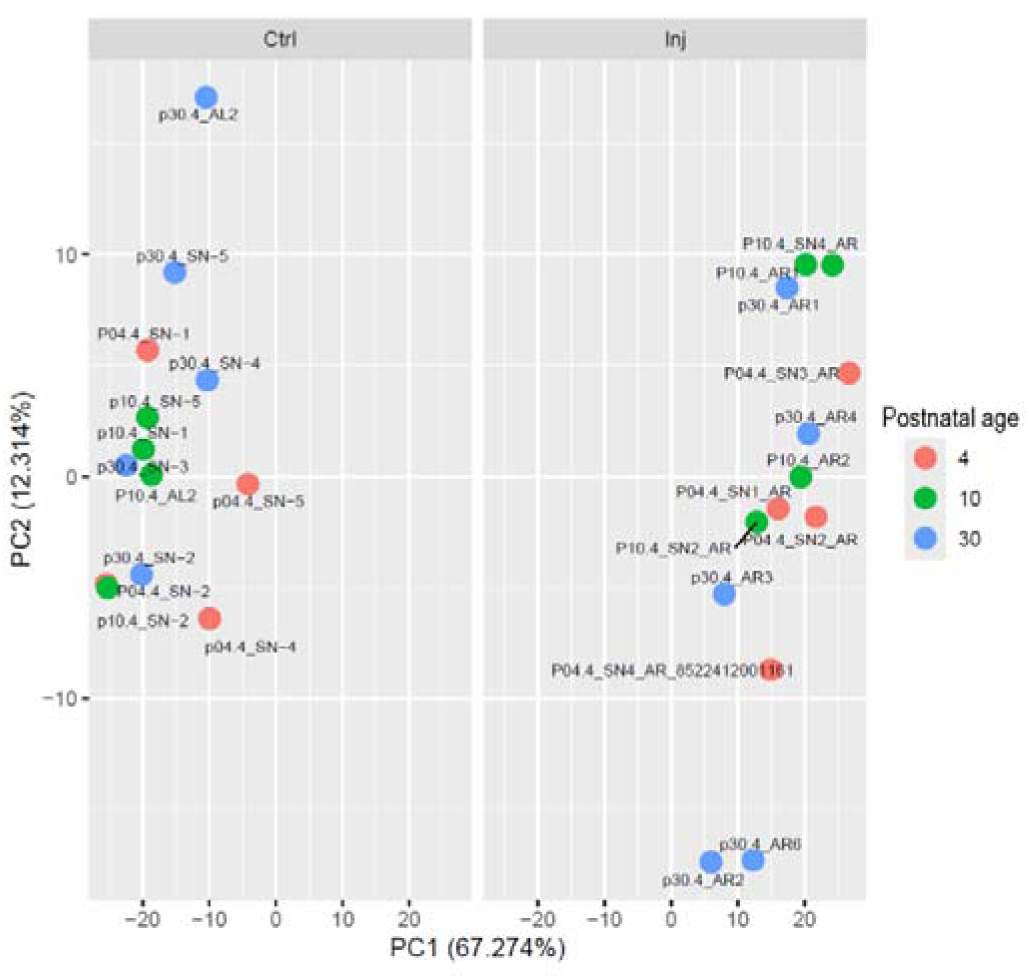
PCA of the sciatic nerve samples included in this study, control (Ctrl) and injured (Inj). Each dot represents an individual sample, showing spatial segregation by condition.

### Pathway enrichment analysis

Transcriptomic analysis of the distal sciatic nerve in P30 injured mice (Figure 11a) revealed strong activation of several signalling pathways and biological processes. The most prominent were cell adhesion molecules, axon guidance, MAPK signalling, and cytokine–receptor interaction. Processes linked to cell cycle, apoptosis, and DNA replication were also enriched, pointing to cellular turnover and injury responses. Immune and inflammatory pathways were prominent, including TNF, NF-κB, Toll-like and NOD-like receptor signalling, chemokine signalling, and IL-17. Additional enriched pathways included sphingolipid metabolism, Hippo signalling, and lysosomal genes. Overall, these findings indicate that the distal nerve responds to injury by activating genes involved in adhesion, axonal remodelling, inflammation, stress responses, and regulation of cell cycle and cell death. In P4 injured mice (Figure 11b), transcriptomic profiling of the distal nerve showed marked enrichment of inflammatory and immune-related pathways. The most significant were chemokine signalling, sphingolipid metabolism, cell adhesion molecules, and NF-κB signalling. Other enriched pathways included cytokine–receptor interaction, IL-17 signalling, and NOD-like receptor signalling, together with endocytosis and phagosome formation, reflecting immune activation and clearance processes. Further pathways, such as TNF signalling, axon guidance, Toll-like receptors, leukocyte migration, MAPK, Hippo, apoptosis, gap junctions, and RIG-I-like receptor signalling, were also evident. This pattern suggests that at P4, the distal nerve mounts a strong inflammatory, immune, and cellular remodelling response. In P10 injured mice (Figure 11c), a different enrichment profile was observed. The most significant pathways were cell adhesion molecules and focal adhesion, with strong enrichment in axon guidance and actin cytoskeleton regulation. Signalling routes linked to cellular communication and growth, such as Rap1, MAPK, PI3K–Akt, and ErbB, were also prominent. Other enriched processes included leukocyte migration, ECM–receptor interaction, NK cell-mediated cytotoxicity, and fatty acid biosynthesis and elongation. These findings indicate that at P10, the distal nerve activates pathways related to adhesion, axonal and cytoskeletal remodelling, immune cell migration, and metabolism, reflecting both tissue reorganisation and immune responses.

**Figure 11.**
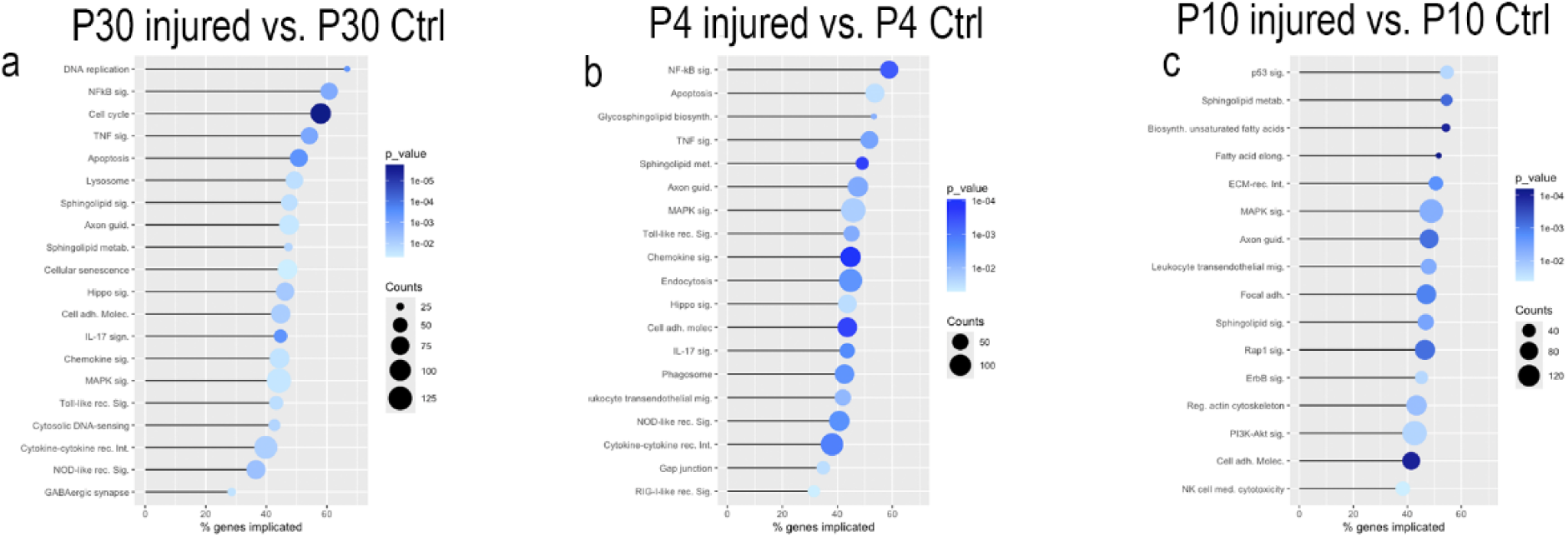
Differential enrichment of KEGG signalling pathways across postnatal stages. Lolliplots display the main enriched signalling pathways identified by gene over-representation analysis ordered by the percentage of genes implicated for that pathway, with p value represented by a colour scale, and number of genes (counts) by a size scale, in sciatic nerve after injury, compared to controls. (a) P30 injured vs. P30 control (b) P4 injured vs. P4 control (c) P10 injured vs. P10 control.

### Gene Set Enrichment Analysis (GSEA)

We performed a customized list of canonical myelin-associated genes (J. Li et al. 2013; Yi et al. 2015; Zhao et al. 2022) , including Mag (myelin-associated glycoprotein), Mal (myelin and lymphocyte protein), Mbp (myelin basic protein), Plp (proteolipid protein), Pmp22 (peripheral myelin protein 22), and Prx (periaxin) to evaluate their expression after injury. For all the selected genes, we observed a negative Log2fold change values following injury. GSEA of these myelination markers revealed significant downregulation across all examined age groups (Figure 12a, Table 1), with negative normalized enrichment scores (NES) across all time points: P4 (NES=-1,928, p.adj=1,86x10^-6^), P10 (NES=-1,978, p.adj=3,75x10^-7^), and P30 (NES=-1,911, p.adj=1,73x10^-5^), indicating consistent and significant downregulation of this gene set following nerve injury.

**Figure 12.**
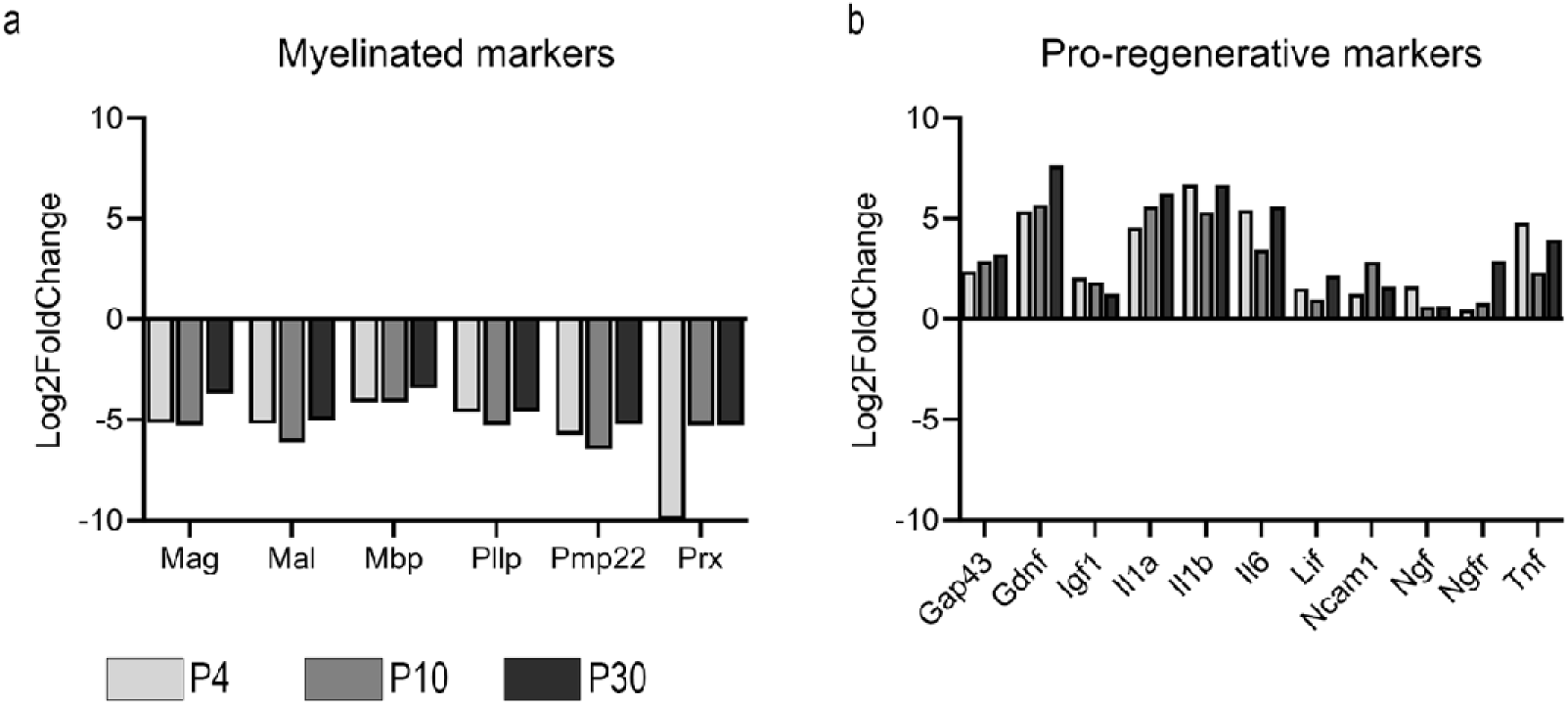
Gene Set Enrichment Analysis (GSEA) showing Log2FoldChange of (a) myelinated markers (Mag, Mal, Mbp, Plp, Pmp22, Prx) and (b) pro-regenerative markers (Gap43, Gdnf, Igf1, Il1a, Il1b, Il6, Lif, Ncam1, Ngf, Ngfr, Tnf) across P4, P10 and P30 injured nerves compared to control ones.

**Figure 13.**
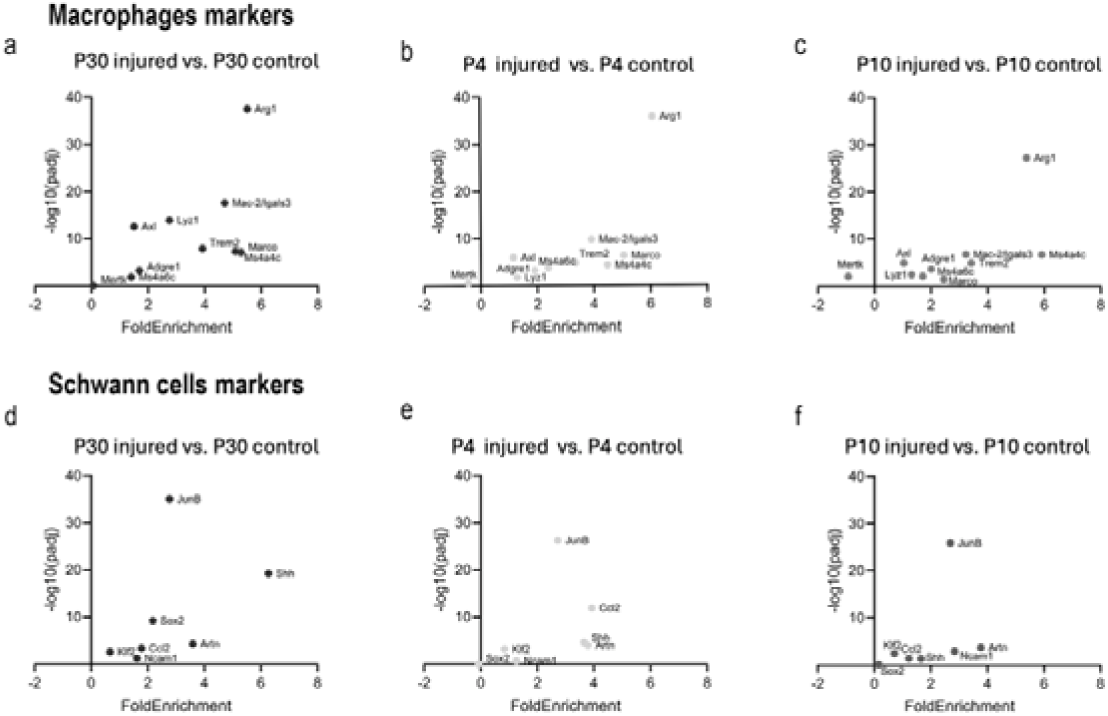
Enrichment of macrophage and Schwann cell marker genes in distal sciatic nerve after transection at different developmental stages. Scatterplots show the fold enrichment and statistical significance (-log10 adjusted p value) for canonical marker genes of macrophages (a-c) and Schwann cells (d-f). Comparisons are shown between injured and control at three ages: (a, d) P30 injured vs. P30 control. (b, e) P4 injured vs. P4 control. (c, f) P10 injured vs. P10 control.

**Table 1.** Results of the Gene Set Enrichment Analysis for mature myelinating Schwann cell markers and pro-regenerative markers in P4, P10, and P30 injured sciatic nerves from mice. Reported values include the enrichment score, NES (normalized enrichment score), and significance values (adjusted P value). A negative NES indicates that the corresponding gene set is downregulated compared with control nerves, whereas a positive NES indicates that the corresponding gene set is upregulated compared to control nerves.

|  | Mature Myelinated SC markers |  |  | Pro-regenerative markers |  |  |
| --- | --- | --- | --- | --- | --- | --- |
|  | P4 | P10 | P30 | P4 | P10 | P30 |
| Enrichment score | -0,977 | -0,976 | -0,972 | 0,757 | 0,675 | 0,774 |
| NES | -1,928 | -1,978 | -1,911 | 1,775 | 1,585 | 1,772 |
| P. adjusted | 1,86E-06 | 3,75E-07 | 1,73E-05 | 0,005 | 0,041 | 0,003 |

We also assessed classical pro-regenerative genes upregulated by SC and macrophages in the distal degenerating stump (Terenghi 1999; Yi et al. 2015; Lee and Cho 2021; Contreras et al. 2022), such as Gap43 (Growth associated protein 43), Gdnf (Glial cell line-derived neurotrophic factor), Igf1 (Insulin-like Growth factor 1), Il1a (Interleukin 1 alpha), Il1b (Interleukin 1 beta), Il6 (Interleukin 6), Lif (Leukemia inhibitory factor), Ncam1 (Neural cell adhesion molecule 1), Ngf (Nerve growth factor), Ngfr (Nerve growth factor receptor) and Tnf (Tumor necrosis factor). These genes displayed similar expression profiles across age groups (Figure 12b, Table 1). GSEA analysis revealed significant enrichment of these markers with positive NES values: P4 (NES=1,775, p.adj=0,005), P10 (NES=1,585, p.adj=0,041), and P30 (NES=1,772, p.adj=0.003).

The enrichment analysis demonstrated statistically significant patterns for both mature myelinating Schwann cell markers and pro-regenerative markers across all age groups, with all adjusted p-values<0.05. The enrichment scores indicated consistent downregulation of myelination programs and upregulation of regenerative programs following sciatic nerve transection across the examined developmental stages.

Analysis of canonical marker gene enrichment revealed dynamic regulation of immune and glial cell populations in the distal sciatic nerve across postnatal development. Macrophage markers showed consistent upregulation at all ages after injury. Genes such as Arg1, Mac-2/Lgals3, Ms4a4c, Trem2, and Marco displayed robust fold enrichment and strong statistical significance (high –log10(padj)), especially at P30 and P4, indicating pronounced macrophage activation in the injured nerve. Notably, Arg1 exhibited the highest enrichment and significance throughout, suggesting strong polarization/activation of macrophages. By contrast, markers like Mertk displayed minimal or negative fold enrichment, indicating selective involvement of subpopulations. Schwann cell markers demonstrated heterogeneous enrichment patterns. Major developmental regulators such as JunB and Ccl2 showed highly significant enrichment after injury at all ages, particularly at P30 and P4. Other markers, including Shh and Artn, were consistently upregulated, while markers like Sox2 and Ncam1 showed modest or variable enrichment, especially at P4 and P10. These data reflect dynamic activation alongside diversity in Schwann cell responses after nerve injury, dependent on age and marker specificity.

## Discussion

It is generally assumed that younger neurons display a greater regenerative capacity following peripheral nerve injury. This notion is mainly based on studies reporting faster regeneration and functional recovery in younger animals (Belin et al., 1996, Hess et al., 2006). However, part of this apparent advantage might be biased by the massive motoneuron death observed in the earliest postnatal stages (Navarrete and Vrbová 1984; Lowrie and Vrbová 1984). This high mortality rate has also limited the number of studies performed in early postnatal stages. Therefore, it is critical to simultaneously assess both neuronal survival and regenerative capacity during these periods.

In this study, we quantified motoneuron survival and regenerative capacity after peripheral nerve injury using histological and molecular approaches, taking advantage of Chat-Cre/Ai9 and Chat-Cre/Ribotag mice models. In addition, we complemented these data with a descriptive and molecular analysis of Wallerian degeneration. All experiments were performed in animals injured at postnatal days 4, 10 and 30 (P4, P10 and P30).

Death and axonal regeneration of motoneurons when injured at different postnatal stages Our results first show that nerve injury at P4 induces a substantial loss of motoneurons (∼50%), whereas no significant neuronal death was observed at P10 or P30. These findings are consistent with previous studies in postnatal rats and mice, that describes a well-defined period of motoneuron loss (P0–P5 in rodents) after nerve lesions (Navarrete and Vrbová 1984; Schmalbruch 1984; Lowrie, Krishnan, and Vrbová 1987; Kemp et al. 2015b). We also explored the regenerative capability of motoneurons when injured at these different postnatal stages. For a fair comparison between postnatal ages, it was important to characterize intact nerves. The number of Chat+ axons observed in the nerve increased with age, fact that can be related to a progressive myelination of axons. Thin and still not myelinated axons may be not detected in the confocal microscope and thus, lower number of fluorescent axons are observed at early postnatal stages compared to juvenile ones. Therefore, for a fair comparison of the regenerative capability at different ages, we normalized the number of regenerating axons by the number of axons in the corresponding contralateral intact nerve. Moreover, the amount of motoneuron death would also impact on the total number of regenerating axons through the distal stump. Therefore, the percentage of regenerating axons after injury was corrected for neuronal death. This correction can have some limitations, since regenerating axons usually extends some sprout when regenerating (Wiezl et al., 2005) and therefore, age differences in extending collaterals could affect this ratio, but still contributes to a better understanding of the regenerative capacity of neurons at different postnatal stage. By doing so, we observed that animals injured at P4 exhibited the highest proportion of regenerating axons, whereas regeneration in P10 and P30 animals did not reach control levels at 14 days post injury. These results support the hypothesis that immature animals possess a strong regenerative potential, which may be masked by the high motoneuron death.

Transcriptional programs activated in axotomized motoneurons at different ages To determine whether these differences depend on the intrinsic growth potential of motoneurons, we analysed the translatome of injured motoneurons at P4, P10 and P30 using the Ribotag assay (Sanz et al. 2019). In juvenile (P30) injured motoneurons, we detected the activation of a regenerative programme, extensively described in the literature (reviewed in Allodi et al., 2012, He and Jin 2016), with overexpression of several regeneration-associated genes (RAGs), including Atf3, Gap43, Ngfr and Ccl2. In contrast, upregulation of classical RAGs was absent in P4 and P10. Despite the lack of upregulation of RAGs, these motoneurons were able to regenerate, fact that suggests that these neurons retain a strong outgrowth potential and, therefore, do not need to switch to a pro-regenerative transcriptional program. However, we cannot rule out that the absence of changes in our dataset may be influenced by the timing of analysis (7 dpi), and earlier timepoints (2–4 dpi) might reveal a more pronounced transcriptional response.

The analysis of specific pathways enriched after injuring juvenile motoneuron supports a mature regenerative programme capable of integrating cell survival mechanisms, metabolic adaptations and tissue remodelling. Pathways related with axonal regeneration and cell survival, such as Hypoxia-related responses (Cho et al., 2015; Smaila et al., 2020) and enrichment in steroid biosynthesis and cholesterol homeostasis (Porcu et al., 2016; Lin et al., 2020; Tang 2022), were upregulated. Inflammatory pathways, including TNF-α/NF-kB, were also strongly upregulated; NF-kB in particular plays a central role in both inflammation and neuronal survival, regulating genes downstream of TNF-α and IL-1β (Anilkumar and Wright-Jin 2024). Other relevant pathways included IL6/Stat3, essential for axonal regrowth (Cafferty et al., 2004), glial activation, proliferation and anti-apoptotic responses (Shimizu, Kiyooka, and Ohshima 2021; Leibinger et al., 2021), and the Neuregulin-1/ErbB system, which supports both axonal regeneration and motoneuron protection under excitotoxic stress (Chang et al., 2013; Mòdol-Caballero et al., 2018; Romero-Ortega et al., 2023). Finally, enrichment of cytoskeletal remodelling and cell migration pathways was observed, consistent with their essential roles in growth cone formation and axonal regrowth (Blanquie and Bradke 2018; Leite, Pinto-Costa, and Sousa 2021). Despite the limited death of injured motoneurons observed at these stages, we found enrichment of apoptosis- and death receptor-related pathways, including the activation of genes such as caspase-3 and oxidative stress responses (Martin, Chen, and Liu 2005).

In terms of signalling, surviving motoneurons at P4 displayed enrichment in pathways linked to elevated ATP production, notably oxidative phosphorylation, key element in the aerobic metabolism, that predominates in mature neurons and confers them protection against stressors (Jimenez-Blasco et al., 2024). Foxo1, a key regulator of cell survival under adverse conditions (Du and Zheng 2021) was also upregulated. However, this upregulation seems insufficient to rescue axotomized neurons from death at these stages. Additional enriched pathways included cholesterol homeostasis, relevant for axonal growth (Tang 2022), and estrogenic responses, consistent with reported neuroprotective effects (Fargo et al., 2009). Classical pathways related with death were not upregulated despite the marked motoneuron death detected at this stage.

In P10 animals, no significant motoneuron loss was observed compared to uninjured controls, but regenerative capacity was not as good as in P4 and P30 animals. Previous studies have reported that regeneration is limited at this developmental window (P10–P15), likely due to progressive loss of plasticity, increased neurotrophic dependence and a more pro-inflammatory environment (Fitzgerald and McKelvey 2016). A down-regulation of some RAGs after the injury could contribute to this impaired regenerative capability. Moreover, the increase of T3 hormone levels during this period have been related with a reduced axonal regenerative potential (Avci et al., 2012). The combination of an unfavourable environment and reduced growth factor signalling could explain the poor RAG upregulation (Schwab and Bartholdi 1996; Oishi et al., 2004). The signalling profile at P10 was markedly different from P4 and P30. Specifically, we found widespread downregulation of mTORC1/TOR, a central pathway for axonal growth and regeneration (Switon et al., 2017). Inhibition of cytoskeletal remodelling and axonal transport pathways, processes essential for growth cone dynamics, were also observed (Bisby and Tetzlaff 1992; Blanquie and Bradke 2018). Downregulation of cellular respiration pathways could lead to an insufficient energy production to support regeneration. Finally, autophagy response was downregulated, which is critical for organelle clearance and cellular renewal (Huang et al., 2016; Chen, Deng, and Zhang 2025), further contributing to reduced regenerative competence.

### Wallerian degeneration of the distal injured nerve at different postnatal stages

Since we evaluated the regenerative capability of motoneurons in vivo, extrinsic factors must also be considered, particularly Wallerian degeneration. Axonal regeneration depends largely on the efficiency of this process, which clears myelin and axonal debris to create a permissive environment (Rotshenker 2011). To characterise this response, we used a complete sciatic nerve transection model without repair, which prevents spontaneous regeneration and allows a precise description of degeneration dynamics during the first post-injury week.

In P30 and P10 mice, nerve injury triggers the canonical degenerative cascade, characterised by progressive myelin breakdown, lipid accumulation, macrophage recruitment and the transition of Schwann cells towards a repair phenotype (Geuna et al., 2009; Gaudet, Popovich, and Ramer 2011) as previously described (Rotshenker 2011). In contrast, P4 nerves degenerated rapidly, with near-complete myelin loss within 2–4 dpi, minimal increase of macrophages (CD68), low transient lipid accumulation and scarce myelin remnants (LFB).

Despite some differences at the molecular level, at all ages studied we could observe a successful downregulation of Schwann cell myelination genes, such as Mag, Mal, Mbp, Pllp, Pmp22 and Prx (J. Li et al., 2013; Yi et al., 2015; Zhao et al., 2022). and an up regulation of pro-regenerative ones such as Gap43, Gdnf, Igf1a, Il1a, Il1b, Lif, Ncam1, Ngf, Ngfr and Tnf (Terenghi 1999; Yi et al., 2015; Lee and Cho 2021; Contreras et al., 2022). It is worth to note that, despite the reduced number of macrophages observed in degenerating nerves injured at P4, upregulation of genes whose products are secreted by macrophages during Wallerian degeneration are also observed at this stage. Again, it seems that the low quantity of these cells in the degenerating nerves of these animals does not impact on the pro-regenerative environment of the distal nerve. The apparent contradiction of our findings at P4 (low increase of macrophages in the injured nerve but an inflammatory profile like the one observed at P30) could be linked with the lower amount of myelin observed in immature nerves. In these circumstances, a modest increase of macrophages is sufficient to phagocyte the myelin, that in fact is rapidly cleared; but these macrophages, together with Schwann cells, orchestrate a good pro-regenerative response, that contribute to the successful growth of the axons of surviving motoneurons. In fact, at all three stages, we also detected strong expression of phagocytic and repair macrophage markers, including Arg1 (Ydens et al., 2012; Stratton et al., 2020; Msheik et al., 2022), Lgals3 (Stratton et al., 2020), Trem2 (Ydens et al., 2012; Stratton et al., 2020; Peshoff et al., 2023) and Marco (Stratton et al., 2020). We also identified markers of repair Schwann cells, such as JunB (Jessen and Mirsky 2016), Ccl2, essential for macrophage recruitment, Shh, which promotes c-Jun activation, and Artn, that supports neuronal survival and axonal regeneration (Jessen and Mirsky 2016; Jessen and Mirsky 2019; Kalinski et al., 2020).

Besides these similarities, pathways were differentially enriched depending on the age. In P30 animals, we observed robust activation of inflammatory and immune pathways, including NOD-like receptors (Ydens et al., 2015), IL-17 signalling, crucial for macrophage recruitment (Bombeiro, Fernandes, and Ribot 2024) (Y. Huang et al., 2024), and TNF-α (Alhamdi et al., 2025). In parallel, we detected strong activation of proliferative and repair programmes, consistent with Schwann cell expansion post-injury (Hung et al., 2015; Painter et al., 2014). Signalling analysis highlighted MAPK/ERK (Lindwall Blom, Mårtensson, and Dahlin 2014; Wei et al., 2024; Liu and Zhou 2025), which promotes Schwann cell reprogramming, and the Hippo pathway (Feltri et al., 2021; Wang et al., 2023), implicated in differentiation towards the myelinating phenotype. We also identified upregulation of sphingolipid metabolism, relevant for neuronal signalling (Meyer zu Reckendorf et al., 2020; C. Li et al., 2024), and lysosomal activation, essential for myelin clearance.

In P4 animals, transcriptomic profiles showed enrichment of chemokine signalling pathways, a hallmark of the neonatal inflammatory response (F. Wang et al., 2025). NF-kB was also activated, driving the production of inflammatory mediators such as TNF-α and IL-1β (Zhang et al., 2025). Despite these differences, P4 shared with P30 the activation of IL-17, MAPK/ERK, Hippo and sphingolipid metabolism, indicating that both ages engage similar degenerative mechanisms to create a permissive environment for regeneration.

By contrast, P10 animals exhibited a transcriptional profile that could contribute to lower regenerative capacity. Notably, we detected activation of p53 pathway and pro-apoptotic genes such as Bax and inhibition of survival genes (Culmsee and Mattson 2005) (Li et al., 2023). Nevertheless, several pathways were shared with P30, including sphingolipid metabolism, adhesion molecule signalling and axon-related pathways. Therefore, at this stage, there is an active response attempting to promote axonal regeneration, albeit not as complete as in juvenile mice.

Taken together, our results indicate that Wallerian degeneration proceeds efficiently at all the developmental stages analysed, despite some molecular or structural differences that could influence axonal regeneration. A lower amount of macrophage infiltration at early postnatal stages only reflects a reduced demand of phagocytosis due to the low myelination ratios at these ages, that does not interfere with the ability of the distal stump to create a permissive milieu for regeneration. In contrast, we observed robust differences in the transcriptional profile of axotomized motoneurons depending on the age when were injured. Therefore, differences in regenerative capacity seems primarily determined by the intrinsic growth state of motoneurons at the time of injury. At P4, motoneurons remain in a growth state reminiscent of the embryonic stage and therefore do not require the activation of a dedicated pro-regenerative transcriptional programme, as occurs in P30 neurons. In contrast, juvenile motoneurons undergo a marked upregulation of regeneration-associated genes (RAGs), which are essential to ensure axonal regeneration in mature peripheral neurons. P10 neurons, however, could be in a transitional stage: they have already begun to downregulate their growth-related profile but still fail to efficiently activate a pro-regenerative programme after injury. This may underlie a slightly lower regenerative capacity compared with both younger postnatal and more mature neurons. However, it is important to remark that motoneurons at all ages retained a good regenerative ability, and the major limiting factor in early postnatal stages is the massive neuronal death.

## Conclusion

Despite the massive loss observed in motoneurons when axotomized at P4, surviving ones shown a robust regenerative capacity, that surprisingly is not dependent on the activation of RAGs. These neurons probably do not need to switch to a pro-regenerative program since still retaining an embryonic growth like state. On the other hand, although early studies proposed that the immaturity of Schwann cells might contribute to neuronal loss at these early stages, our results indicate that Wallerian degeneration is already efficient in both clearing myelin debris and creating a permissive pro-regenerative environment at neonatal stages. Instead, it is the transcriptomic profile of injured motoneurons that differs substantially with age, highlighting intrinsic neuronal factors as the key determinants of regenerative success. Further investigation into the pathways that are differentially regulated between P4, P10 and P30 motoneurons could therefore provide critical insights into the mechanisms driving motoneuron vulnerability and death at early postnatal stages

## Acknowledgements

The authors appreciate the technical help of Neus Hernández. Mònica Espejo and Jessica Jaramillo.

## Author Contributions

Beatriu Molina Esteve, Natalia Lago, and Esther Udina contributed to the conception and design of the study. Material preparation, data collection, and analysis were performed by all authors (Beatriu Molina Esteve, Marina Pujol-Masip, David Ovelleiro, José Antonio Gómez-Sánchez, Natalia Lago, and Esther Udina). The first draft of the manuscript was written by Beatriu Molina Esteve, and all authors commented on previous versions of the manuscript. Supervision and critical revision for important intellectual content were provided by Natalia Lago and Esther Udina. All authors read and approved the final manuscript.

