## Supplementary data for "Assessment of Motoneuronal Regeneration and Wallerian Degeneration Following Axotomy in Postnatal Mice"

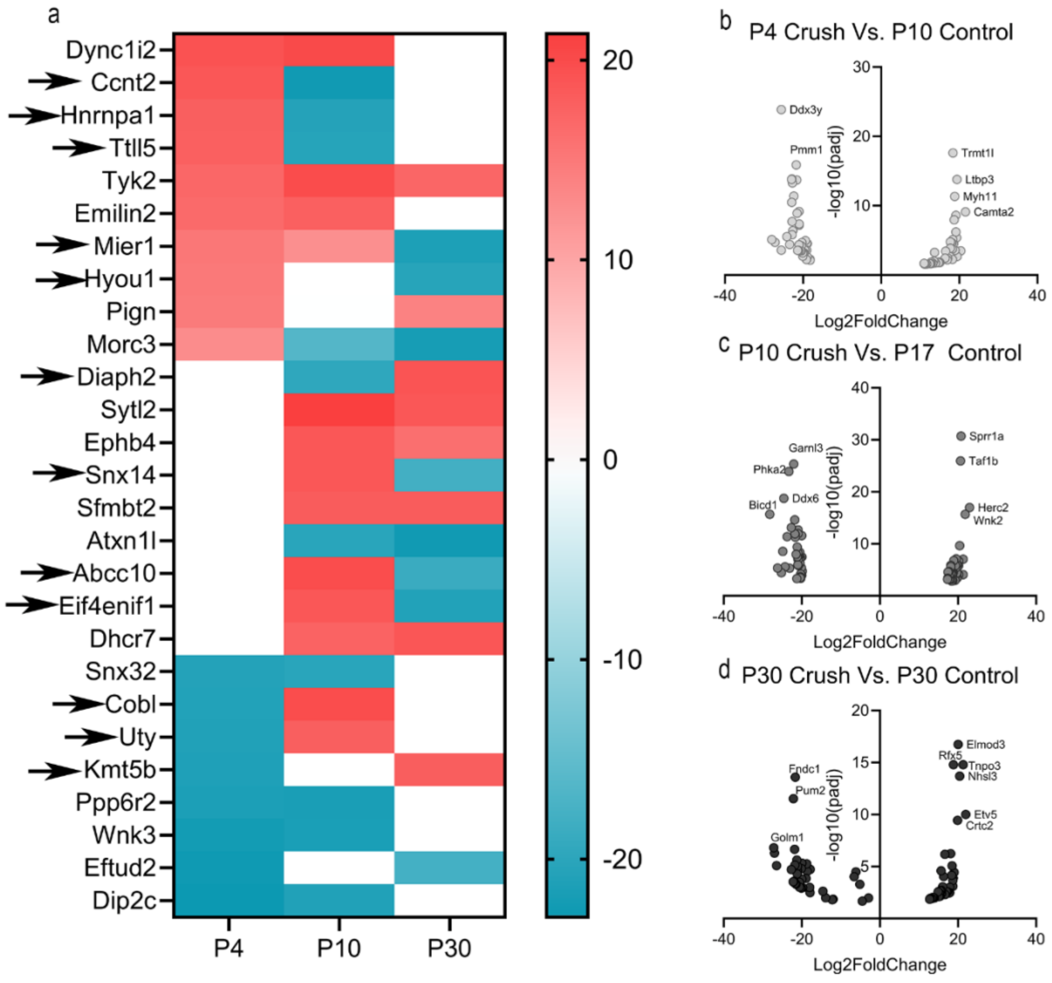


**Supplementary Figure 1**. Changes in gene expression 7 days after the crush injury in the motoneuron translatome in P4, P10 and P30 mice. (a) Heatmap showing the variation in expression of selected genes at P4, P10 and P30. Genes highlighted with arrows were selected for their relevance. The colour scale indicates the range of expression (Log2FoldChange), from upregulation (red) to downregulation (blue). (b-d) Volcano plots showing changes in gene expression (Log2FoldChange) and significance (-log10(padj)) between crush and control conditions for the indicated time point comparisons. (b) P4 Crush vs. P10 Control (c) P10 Crush vs. P17 Control (d) P30 Crush vs. P30 Control. Labelled gene names highlight those most significantly regulated in each contrast.

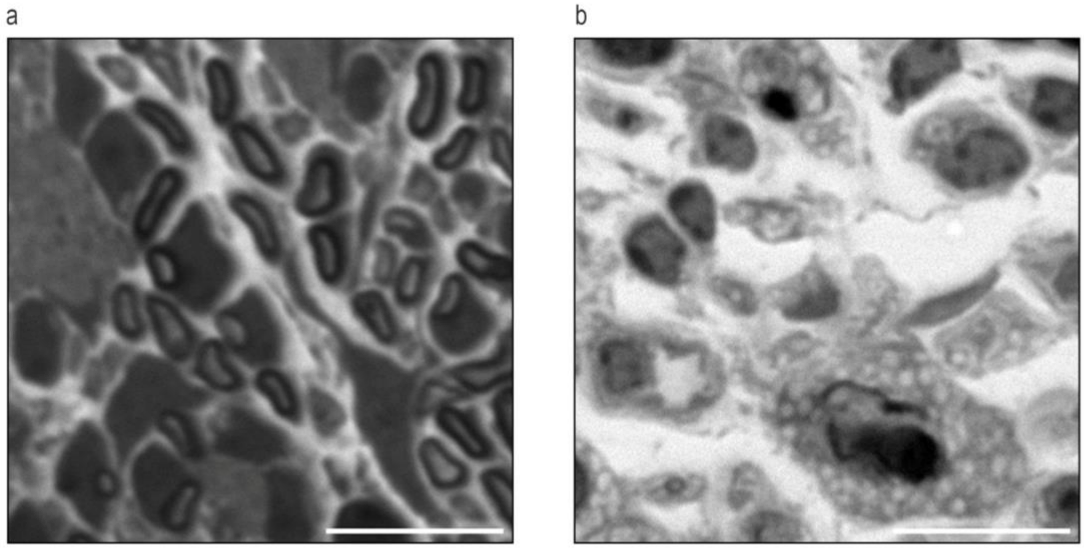


**Supplementary Figure 2**. (a) Representative image of uninjured nerve from P4 mice, showing the immature myelinating nerves. (b) Representative image of phagocyting bodies present after an injury in the sciatic nerve. Scale bar: 10 µm

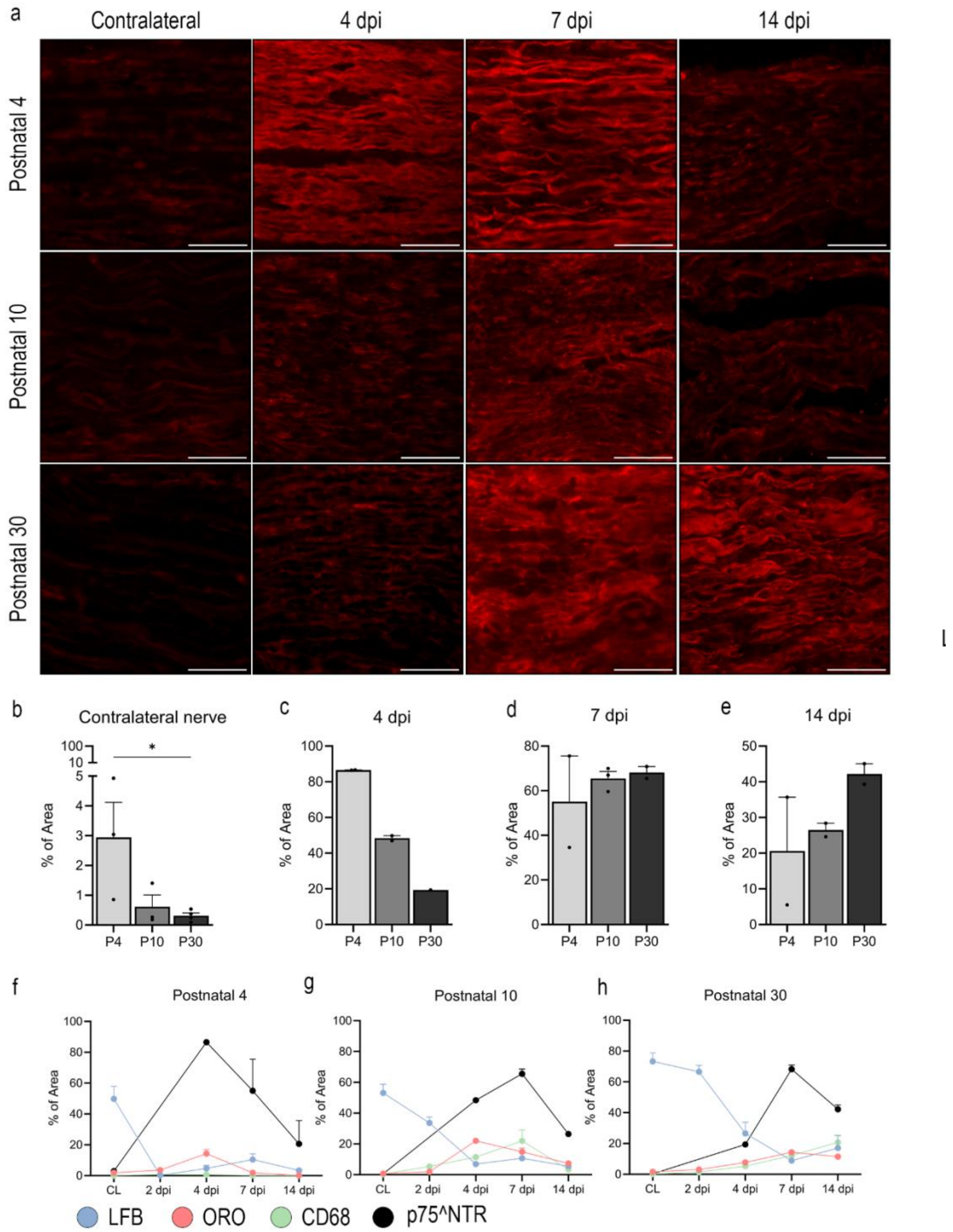


**Supplementary Figure 3**. (a) Representative images of p75^NTR immunostaining in injured nerves of P4, P10 and P30 mice 4,7 and 14 dpi and its contralateral. (b-e) Percentage of area stained with anti-p75^NTR in P4, P10 and P30 contralateral and injured nerves. Statistics: Kruskal-Wallis test. $p<0.05$. (f-g) Percentage of area stained with LFB, ORO, CD68 and p75^NTR of contralateral and injured nerves of P4, P10 and P30 mice 2, 4, 7 and 14 dpi.

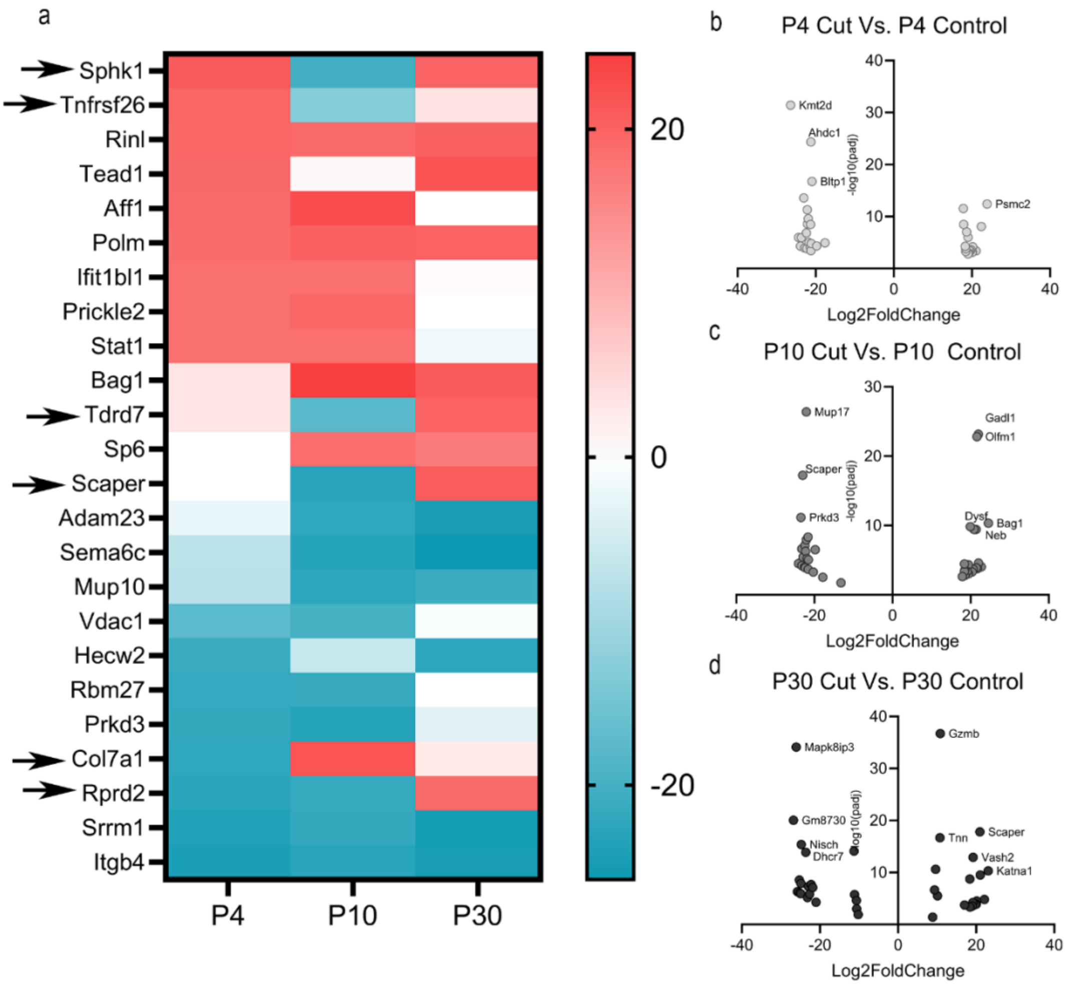


**Supplementary Figure 4**. Changes in gene expression 4 days after the cut injury in the Sciatic nerve transcriptome in P4, P10 and P30 mice. (a) Heatmap showing the variation in expression of selected genes at P4, P10 and P30. Genes highlighted with arrows were selected for their relevance. The colour scale indicates the range of expression (Log2FoldChange), from upregulation (red) to downregulation (blue). (b-d) Volcano plots showing changes in gene expression (Log2FoldChange) and significance (-log10(padj)) between cut and control conditions for the indicated time point comparisons. (b) P4 Cut vs. P4 Control (c) P10 Cut vs. P10 Control (d) P30 Cut vs. P30 Control. Labelled gene names highlight those most significantly regulated in each contrast.

| | P30 | | P4 | | P10 | |
|---|---|---|---|---|---|---|
| Gene ID | Log2Fold Change | padj | Log2Fold Change | padj | Log2Fold Change | padj |
| Katna1 | 23,079 | 4,79E-11 | 1,502 | 7,65E-01 | 0,545 | 9,15E-01 |
| Tead1 | 22,109 | 1,52E-05 | 19,277 | 1,11E-03 | 0,925 | 9,01E-01 |
| Bag1 | 21,044 | 2,99E-10 | 3,407 | 4,31E-01 | 24,565 | 4,58E-11 |

| | | | | | | |
|---|---|---|---|---|---|---|
| Scaper | 20,935 | 1,52E-18 | -0,224 | 9,52E-01 | -23,011 | 5,70E-18 |
| Rinl | 20,145 | 2,93E-05 | 19,521 | 4,20E-04 | 19,132 | 4,30E-04 |
| Sphk1 | 20,040 | 1,10E-04 | 21,050 | 3,90E-04 | -20,271 | 5,10E-04 |
| Polm | 20,036 | 1,10E-04 | 18,986 | 1,51E-03 | 20,358 | 4,80E-04 |
| Sh3pxd2 | 19,945 | 3,64E-05 | -14,824 | 1,04E-02 | 0,256 | 9,72E-01 |
| Tdrd7 | 19,824 | 1,40E-04 | 3,307 | 6,41E-01 | -17,839 | 2,92E-03 |
| Vash2 | 19,223 | 1,22E-13 | 0,803 | 8,28E-01 | 1,053 | 7,68E-01 |
| Brpf1 | 19,143 | 6,29E-05 | 0,234 | 9,77E-01 | 2,619 | 6,80E-01 |
| Rprd2 | 18,984 | 2,70E-04 | -22,826 | 1,10E-04 | -21,585 | 2,00E-04 |
| Eif2b4 | 18,433 | 4,30E-04 | -5,007 | 4,69E-01 | 0,352 | 9,64E-01 |
| Shf | 18,362 | 1,74E-09 | -0,617 | 8,94E-01 | -0,879 | 8,37E-01 |
| Sp6 | 17,042 | 1,70E-04 | -0,165 | 9,82E-01 | 18,526 | 2,60E-04 |
| Gzmb | 10,827 | 1,92E-37 | 8,094 | 3,96E-21 | 6,473 | 1,05E-09 |
| Tnn | 10,735 | 2,07E-17 | 7,651 | 1,73E-07 | 10,393 | 2,86E-13 |
| Gria1 | 10,126 | 3,03E-06 | 8,451 | 7,80E-04 | 2,823 | 2,42E-01 |
| Cxcl3 | 9,612 | 2,47E-11 | 9,266 | 1,65E-08 | 7,048 | 2,00E-05 |
| Myh3 | 9,336 | 2,30E-07 | 6,456 | 1,80E-03 | 6,239 | 2,09E-03 |
| Ankrd1 | 7,843 | 4,00E-04 | 21,384 | 1,54E-16 | 7,898 | 1,73E-03 |
| Atp1b4 | 6,703 | 1,02E-03 | 3,122 | 1,47E-01 | 10,997 | 7,11E-07 |
| Igfn1 | 5,433 | 7,00E-04 | 2,085 | 3,14E-01 | 11,143 | 2,39E-09 |
| Tnfrsf26 | 3,597 | 5,33E-01 | 19,798 | 2,50E-04 | -13,156 | 1,84E-02 |

| | | | | | | |
|---|---|---|---|---|---|---|
| Col7a1 | 2,713 | 6,79E-01 | -22,292 | 1,40E-04 | 21,751 | 1,60E-04 |
| Ahdc1 | 2,655 | 1,41E-01 | -21,227 | 4,53E-25 | 1,657 | 4,24E-01 |
| Tro | 2,094 | 7,44E-01 | 0,519 | 9,47E-01 | 20,639 | 1,80E-04 |
| Nfia | 2,017 | 5,41E-01 | -0,816 | 8,36E-01 | -22,784 | 1,06E-13 |
| Adcy7 | 2,014 | 5,90E-01 | 0,462 | 9,21E-01 | 21,396 | 3,58E-10 |
| Fndc3b | 1,807 | 7,56E-01 | 20,197 | 6,62E-05 | -0,301 | 9,64E-01 |
| Slc6a2 | 1,522 | 7,91E-01 | -21,122 | 1,52E-05 | 0,235 | 9,72E-01 |
| C5ar2 | 1,518 | 6,95E-01 | 18,625 | 8,74E-08 | 0,871 | 8,40E-01 |
| Aftph | 1,440 | 6,62E-01 | 17,880 | 3,41E-09 | 1,124 | 7,52E-01 |
| Dysf | 1,244 | 7,24E-01 | 0,932 | 8,12E-01 | 19,939 | 1,47E-10 |
| Slc25a14 | 1,027 | 8,63E-01 | -1,570 | 8,00E-01 | -21,626 | 7,72E-06 |
| Olfm1 | 1,026 | 6,49E-01 | 2,290 | 3,09E-01 | 21,640 | 1,55E-23 |
| Wdr26 | 0,658 | 9,31E-01 | 20,065 | 7,40E-04 | 0,082 | 9,92E-01 |
| Gadl1 | 0,606 | 8,03E-01 | 2,272 | 3,22E-01 | 21,978 | 7,04E-24 |
| Ifit1bl1 | 0,472 | 9,44E-01 | 18,442 | 5,10E-04 | 18,434 | 4,00E-04 |
| Kmt2a | 0,421 | 9,51E-01 | -13,267 | 1,93E-02 | 22,036 | 2,62E-05 |
| Phlpp2 | 0,150 | 9,84E-01 | -5,356 | 4,44E-01 | -24,075 | 3,16E-05 |
| Uvssa | 0,132 | 9,81E-01 | -22,381 | 1,41E-07 | -1,198 | 8,22E-01 |
| Ankrd2 | 0,128 | 9,79E-01 | 21,438 | 2,31E-09 | -0,057 | 9,91E-01 |
| Ryr1 | 0,096 | 9,90E-01 | 17,263 | 4,22E-03 | 17,890 | 2,36E-03 |
| Ilf3 | 0,080 | 9,87E-01 | 22,438 | 8,22E-09 | 1,945 | 6,71E-01 |

| | | | | | | |
|---|---|---|---|---|---|---|
| Rbm27 | -0,009 | 9,99E-01 | -21,748 | 9,48E-06 | -21,548 | 8,66E-06 |
| Aff1 | -0,037 | 9,96E-01 | 19,105 | 1,40E-03 | 22,723 | 9,06E-05 |
| Primpol | -0,041 | 9,94E-01 | -0,320 | 9,57E-01 | -22,414 | 7,31E-08 |
| Zbtb43 | -0,074 | 9,79E-01 | 21,594 | 5,24E-22 | 0,382 | 8,86E-01 |
| Prickle2 | -0,149 | 9,81E-01 | 18,409 | 1,50E-04 | 19,430 | 4,56E-05 |
| Neb | -0,321 | 9,41E-01 | -0,303 | 9,49E-01 | 20,854 | 3,69E-10 |
| Tyrp1 | -0,625 | 9,35E-01 | -6,402 | 3,39E-01 | 18,551 | 1,56E-03 |
| Spg7 | -0,852 | 8,79E-01 | -2,047 | 7,16E-01 | -23,220 | 2,35E-07 |
| Ttbk2 | -0,856 | 8,21E-01 | -22,172 | 4,79E-12 | -1,519 | 6,85E-01 |
| Nlgn3 | -0,955 | 7,86E-01 | -23,026 | 2,52E-14 | -1,618 | 6,42E-01 |
| Rnf213 | -0,965 | 7,39E-01 | -3,248 | 2,39E-01 | -24,922 | 4,33E-22 |
| Pcbd2 | -1,097 | 8,74E-01 | 0,350 | 9,65E-01 | 21,431 | 1,20E-04 |
| Scn8a | -1,109 | 6,84E-01 | 17,804 | 2,90E-12 | -1,668 | 5,45E-01 |
| Kmt2d | -1,174 | 6,08E-01 | -26,407 | 3,95E-32 | -0,765 | 7,64E-01 |
| Bltp1 | -1,390 | 5,87E-01 | -20,970 | 1,77E-17 | -3,685 | 1,34E-01 |
| Csrnp1 | -1,600 | 7,19E-01 | 0,720 | 8,92E-01 | -22,058 | 1,14E-08 |
| Psmc2 | -1,613 | 6,59E-01 | 23,877 | 3,81E-13 | -0,771 | 8,51E-01 |
| Slc25a13 | -1,887 | 6,47E-01 | -22,120 | 2,53E-09 | -2,904 | 4,85E-01 |
| Stat1 | -1,975 | 6,96E-01 | 18,292 | 5,18E-05 | 18,421 | 3,41E-05 |
| Mup17 | -2,102 | 2,63E-01 | 16,566 | 4,04E-15 | -22,047 | 4,18E-27 |
| Celf1 | -2,571 | 6,99E-01 | 20,148 | 7,10E-04 | -0,199 | 9,80E-01 |

| | | | | | | |
|---|---|---|---|---|---|---|
| Tmod2 | -3,021 | 4,17E-01 | -21,312 | 3,34E-09 | -2,989 | 4,55E-01 |
| Prkd3 | -3,316 | 3,44E-01 | -21,896 | 2,65E-10 | -23,455 | 6,91E-12 |
| Ppfia3 | -3,619 | 3,39E-01 | -3,558 | 4,01E-01 | -21,601 | 4,69E-09 |
| Cobl | -3,883 | 4,03E-01 | -2,742 | 6,11E-01 | -22,227 | 5,24E-07 |
| Nlrx1 | -6,673 | 7,59E-02 | 18,987 | 9,96E-07 | 1,003 | 8,35E-01 |
| Ppp6r2 | -7,424 | 2,05E-01 | -21,210 | 3,50E-04 | 0,718 | 9,24E-01 |
| Smtn | -10,212 | 1,20E-02 | -6,555 | 1,78E-01 | -4,593 | 3,39E-01 |
| Ttc17 | -10,614 | 9,00E-04 | -9,494 | 1,04E-02 | -3,992 | 2,85E-01 |
| Rps3a3 | -10,651 | 2,40E-05 | 17,156 | 1,59E-09 | -0,063 | 9,87E-01 |
| Rps3a2 | -11,148 | 1,77E-06 | 2,373 | 4,34E-01 | -0,128 | 9,70E-01 |
| Cntn1 | -11,287 | 7,85E-15 | -5,922 | 5,64E-06 | -7,233 | 2,93E-08 |
| Mup10 | -21,015 | 5,10E-05 | -7,639 | 2,53E-01 | -22,577 | 1,00E-04 |
| Mup10 | -21,015 | 5,10E-05 | -7,639 | 2,53E-01 | -22,577 | 1,00E-04 |
| Ptk2b | -21,824 | 7,78E-08 | -8,419 | 9,35E-02 | 1,613 | 7,76E-01 |
| Hecw2 | -22,640 | 1,44E-06 | -21,337 | 7,87E-05 | -6,297 | 2,86E-01 |
| Hecw2 | -22,640 | 1,44E-06 | -21,337 | 7,87E-05 | -6,297 | 2,86E-01 |
| Epha4 | -22,959 | 5,01E-08 | -1,060 | 8,68E-01 | 0,735 | 9,05E-01 |
| Ino80d | -23,182 | 6,71E-06 | -0,363 | 9,65E-01 | 17,699 | 2,60E-03 |
| Ino80d | -23,182 | 6,71E-06 | -0,363 | 9,65E-01 | 17,699 | 2,65E-03 |
| Dhcr7 | -23,638 | 1,36E-14 | -5,332 | 1,68E-01 | -3,301 | 3,96E-01 |
| Nisch | -24,799 | 4,12E-16 | -2,898 | 4,68E-01 | 0,987 | 8,19E-01 |

| | | | | | | |
|---|---|---|---|---|---|---|
| Itgb4 | -24,878 | 1,29E-08 | -24,375 | 1,07E-06 | -22,926 | 3,71E-06 |
| Itgb4 | -24,878 | 1,29E-08 | -24,375 | 1,07E-06 | -22,926 | 3,71E-06 |
| Adam23 | -24,892 | 1,18E-06 | -2,406 | 7,43E-01 | -22,339 | 1,20E-04 |
| Adam23 | -24,892 | 1,18E-06 | -2,406 | 7,43E-01 | -22,339 | 1,20E-04 |
| Srp54c | -25,241 | 8,13E-07 | -16,810 | 6,32E-03 | -2,287 | 7,47E-01 |
| Srrm1 | -25,260 | 2,80E-09 | -23,679 | 1,16E-06 | -21,889 | 5,88E-06 |
| Srrm1 | -25,260 | 2,80E-09 | -23,679 | 1,16E-06 | -21,889 | 5,88E-06 |
| Sema6c | -25,766 | 4,67E-07 | -7,251 | 2,80E-01 | -23,209 | 6,22E-05 |
| Sema6c | -25,766 | 4,67E-07 | -7,251 | 2,80E-01 | -23,209 | 6,22E-05 |
| Mapk8ip3 | -26,047 | 7,42E-35 | -3,177 | 2,01E-01 | -2,589 | 2,95E-01 |
| Gm8730 | -26,764 | 8,59E-21 | 16,931 | 3,99E-07 | -1,451 | 7,06E-01 |
| Thsd4 | -30,000 | 6,26E-09 | 2,430 | 7,40E-01 | -0,032 | 9,96E-01 |

**Supplementary table 2.** 120 most up and down regulated genes expressed in the injured motoneurons at P30, P4 and P10 vs uninjured motoneurons at P30, P10 and P17 respectively.

| | P30 | | P4 | | P10 | |
|---|---|---|---|---|---|---|
| Gene ID | Log2Fold Change | padj | Log2Fold Change | padj | Log2Fold Change | padj |
| Katna1 | 23,07917 | 4,79E-11 | 1,501669 | 0,76498 | 0,545466 | 0,91469 |
| Tead1 | 22,10926 | 1,52E-05 | 19,27691 | 0,00111 | 0,924681 | 0,90086 |
| Bag1 | 21,04386 | 2,99E-10 | 3,407312 | 0,43132 | 24,56493 | 4,58E-11 |
| Scaper | 20,93481 | 1,52E-18 | -0,22382 | 0,95179 | -23,0106 | 5,70E-18 |
| Rinl | 20,14543 | 2,93E-05 | 19,52082 | 0,00042 | 19,13224 | 0,00043 |
| Sphk1 | 20,04023 | 0,00011 | 21,0495 | 0,00039 | -20,2712 | 0,00051 |
| Polm | 20,03607 | 0,00011 | 18,98626 | 0,00151 | 20,35802 | 0,00048 |

| | | | | | | |
|---|---|---|---|---|---|---|
| Sh3pxd2 | 19,94495 | 3,64E-05 | -14,824 | 0,01035 | 0,256432 | 0,97216 |
| Tdrd7 | 19,82364 | 0,00014 | 3,307255 | 0,64093 | -17,8393 | 0,00292 |
| Vash2 | 19,22289 | 1,22E-13 | 0,803446 | 0,82777 | 1,052568 | 0,76757 |
| Brpf1 | 19,14345 | 6,29E-05 | 0,233679 | 0,97664 | 2,618925 | 0,68036 |
| Rprd2 | 18,98411 | 0,00027 | -22,8264 | 0,00011 | -21,5846 | 0,00020 |
| Eif2b4 | 18,43305 | 0,00043 | -5,00663 | 0,46853 | 0,352486 | 0,96391 |
| Shf | 18,36205 | 1,74E-09 | -0,61727 | 0,89353 | -0,87867 | 0,83714 |
| Sp6 | 17,04172 | 0,00017 | -0,16495 | 0,98236 | 18,52577 | 0,00026 |
| Gzmb | 10,82656 | 1,92E-37 | 8,093789 | 3,96E-21 | 6,473336 | 1,05E-09 |
| Tnn | 10,73501 | 2,07E-17 | 7,651223 | 1,73E-07 | 10,39341 | 2,86E-13 |
| Gria1 | 10,12552 | 3,03E-06 | 8,451137 | 0,00078 | 2,823461 | 0,24195 |
| Cxcl3 | 9,612205 | 2,47E-11 | 9,265997 | 1,65E-08 | 7,048408 | 2,00E-05 |
| Myh3 | 9,336363 | 2,30E-07 | 6,456478 | 0,00180 | 6,238639 | 0,00209 |
| Ankrd1 | 7,842653 | 0,00040 | 21,38376 | 1,54E-16 | 7,898072 | 0,00173 |
| Atp1b4 | 6,702862 | 0,00102 | 3,122306 | 0,14744 | 10,9968 | 7,11E-07 |
| Igfn1 | 5,432694 | 0,00070 | 2,084802 | 0,31391 | 11,14282 | 2,39E-09 |
| Tnfrsf26 | 3,596619 | 0,5332 | 19,79766 | 0,00025 | -13,1559 | 0,01838 |
| Col7a1 | 2,712504 | 0,67912 | -22,292 | 0,00014 | 21,75071 | 0,00016 |
| Ahdc1 | 2,655241 | 0,14100 | -21,2267 | 4,53E-25 | 1,657074 | 0,42398 |
| Tro | 2,094214 | 0,74429 | 0,519323 | 0,94684 | 20,63874 | 0,00018 |
| Nfia | 2,016632 | 0,54116 | -0,81551 | 0,83628 | -22,7835 | 1,06E-13 |
| Adcy7 | 2,014264 | 0,58960 | 0,462076 | 0,9207 | 21,39553 | 3,58E-10 |
| Fndc3b | 1,807116 | 0,75550 | 20,19653 | 6,62E-05 | -0,30123 | 0,96391 |
| Slc6a2 | 1,521895 | 0,79112 | -21,1218 | 1,52E-05 | 0,234517 | 0,97160 |
| C5ar2 | 1,518206 | 0,6954 | 18,62507 | 8,74E-08 | 0,870767 | 0,83996 |

| | | | | | | |
|---|---|---|---|---|---|---|
| Aftph | 1,440322 | 0,66156 | 17,88008 | 3,41E-09 | 1,123961 | 0,75209 |
| Dysf | 1,243534 | 0,7244 | 0,932058 | 0,81235 | 19,939 | 1,47E-10 |
| Slc25a14 | 1,027367 | 0,86286 | -1,5699 | 0,80000 | -21,6259 | 7,72E-06 |
| Olfm1 | 1,026024 | 0,64894 | 2,290001 | 0,3094 | 21,64016 | 1,55E-23 |
| Wdr26 | 0,65847 | 0,93053 | 20,0654 | 0,00074 | 0,081922 | 0,99199 |
| Gadl1 | 0,606065 | 0,80312 | 2,271605 | 0,32153 | 21,97817 | 7,04E-24 |
| Ifit1bl1 | 0,472155 | 0,94436 | 18,44224 | 0,00051 | 18,43353 | 0,00040 |
| Kmt2a | 0,420976 | 0,9508 | -13,2673 | 0,01930 | 22,03587 | 2,62E-05 |
| Phlpp2 | 0,14996 | 0,98412 | -5,35602 | 0,44377 | -24,075 | 3,16E-05 |
| Uvssa | 0,131878 | 0,98143 | -22,3807 | 1,41E-07 | -1,19752 | 0,82160 |
| Ankrd2 | 0,128102 | 0,97892 | 21,43772 | 2,31E-09 | -0,05719 | 0,99084 |
| Ryr1 | 0,095668 | 0,98979 | 17,26313 | 0,00422 | 17,89004 | 0,00236 |
| Ilf3 | 0,079708 | 0,98726 | 22,43789 | 8,22E-09 | 1,944516 | 0,67076 |
| Rbm27 | -0,0092 | 0,9986 | -21,7482 | 9,48E-06 | -21,5478 | 8,66E-06 |
| Aff1 | -0,03689 | 0,99589 | 19,10513 | 0,00140 | 22,72264 | 9,06E-05 |
| Primpol | -0,04058 | 0,99394 | -0,32045 | 0,95738 | -22,4142 | 7,31E-08 |
| Zbtb43 | -0,07432 | 0,97910 | 21,59444 | 5,24E-22 | 0,382009 | 0,88592 |
| Prickle2 | -0,14856 | 0,98143 | 18,40905 | 0,00015 | 19,4296 | 4,56E-05 |
| Neb | -0,32073 | 0,94060 | -0,30264 | 0,94882 | 20,85406 | 3,69E-10 |
| Tyrp1 | -0,62493 | 0,93518 | -6,40238 | 0,33914 | 18,55115 | 0,00156 |
| Spg7 | -0,85249 | 0,87927 | -2,04719 | 0,71621 | -23,2201 | 2,35E-07 |
| Ttbk2 | -0,85585 | 0,82099 | -22,1718 | 4,79E-12 | -1,51884 | 0,68488 |
| Nlgn3 | -0,95534 | 0,78598 | -23,0259 | 2,52E-14 | -1,61789 | 0,64249 |

| | | | | | | |
|---|---|---|---|---|---|---|
| Rnf213 | -0,96454 | 0,73933 | -3,24808 | 0,23937 | -24,9219 | 4,33E-22 |
| Pcbd2 | -1,09731 | 0,87388 | 0,350297 | 0,96520 | 21,43066 | 0,00012 |
| Scn8a | -1,10858 | 0,68401 | 17,8039 | 2,90E-12 | -1,66828 | 0,54515 |
| Kmt2d | -1,1742 | 0,60839 | -26,4071 | 3,95E-32 | -0,76497 | 0,76419 |
| Bltp1 | -1,39034 | 0,58682 | -20,9697 | 1,77E-17 | -3,68529 | 0,13359 |
| Csrnp1 | -1,60013 | 0,71923 | 0,719605 | 0,89168 | -22,0582 | 1,14E-08 |
| Psmc2 | -1,61345 | 0,65942 | 23,87661 | 3,81E-13 | -0,771 | 0,85076 |
| Slc25a13 | -1,88655 | 0,64654 | -22,1196 | 2,53E-09 | -2,90383 | 0,48526 |
| Stat1 | -1,97476 | 0,69636 | 18,29234 | 5,18E-05 | 18,42128 | 3,41E-05 |
| Mup17 | -2,10152 | 0,26260 | 16,56602 | 4,04E-15 | -22,0473 | 4,18E-27 |
| Celf1 | -2,57076 | 0,69931 | 20,14806 | 0,00071 | -0,19896 | 0,98043 |
| Tmod2 | -3,02097 | 0,41674 | -21,312 | 3,34E-09 | -2,98943 | 0,45499 |
| Prkd3 | -3,3163 | 0,34408 | -21,8959 | 2,65E-10 | -23,4546 | 6,91E-12 |
| Ppfia3 | -3,61871 | 0,33904 | -3,55762 | 0,40052 | -21,6012 | 4,69E-09 |
| Cobl | -3,88251 | 0,40287 | -2,7418 | 0,61119 | -22,2273 | 5,24E-07 |
| Nlrx1 | -6,67339 | 0,07588 | 18,98705 | 9,96E-07 | 1,002712 | 0,83475 |
| Ppp6r2 | -7,42398 | 0,20453 | -21,2099 | 0,00035 | 0,718171 | 0,92403 |
| Smtn | -10,2116 | 0,01195 | -6,55531 | 0,17758 | -4,59315 | 0,33876 |
| Ttc17 | -10,6141 | 0,0009 | -9,4936 | 0,01037 | -3,99158 | 0,28489 |
| Rps3a3 | -10,6506 | 2,40E-05 | 17,15565 | 1,59E-09 | -0,06275 | 0,98687 |
| Rps3a2 | -11,1476 | 1,77E-06 | 2,372527 | 0,43355 | -0,12813 | 0,96961 |
| Cntn1 | -11,2871 | 7,85E-15 | -5,92241 | 5,64E-06 | -7,23335 | 2,93E-08 |
| Mup10 | -21,0146 | 5,10E-05 | -7,63936 | 0,25300 | -22,5772 | 0,00010 |
| Mup10 | -21,0146 | 5,10E-05 | -7,63936 | 0,25300 | -22,5772 | 0,00010 |

| | | | | | | |
|---|---|---|---|---|---|---|
| Ptk2b | -21,8237 | 7,78E-08 | -8,41912 | 0,09349 | 1,612882 | 0,77637 |
| Hecw2 | -22,6395 | 1,44E-06 | -21,3366 | 7,87E-05 | -6,29716 | 0,28622 |
| Hecw2 | -22,6395 | 1,44E-06 | -21,3366 | 7,87E-05 | -6,29716 | 0,28622 |
| Epha4 | -22,9594 | 5,01E-08 | -1,05954 | 0,86818 | 0,735461 | 0,90499 |
| Ino80d | -23,1817 | 6,71E-06 | -0,36314 | 0,96539 | 17,6985 | 0,0026 |
| Ino80d | -23,1817 | 6,71E-06 | -0,36314 | 0,96539 | 17,6985 | 0,00265 |
| Dhcr7 | -23,6383 | 1,36E-14 | -5,33163 | 0,16779 | -3,30098 | 0,39564 |
| Nisch | -24,7994 | 4,12E-16 | -2,89797 | 0,46765 | 0,987142 | 0,81927 |
| Itgb4 | -24,8784 | 1,29E-08 | -24,3747 | 1,07E-06 | -22,9261 | 3,71E-06 |
| Itgb4 | -24,8784 | 1,29E-08 | -24,3747 | 1,07E-06 | -22,9261 | 3,71E-06 |
| Adam23 | -24,8921 | 1,18E-06 | -2,40584 | 0,74277 | -22,3386 | 0,00012 |
| Adam23 | -24,8921 | 1,18E-06 | -2,40584 | 0,74277 | -22,3386 | 0,00012 |
| Srp54c | -25,2409 | 8,13E-07 | -16,8098 | 0,00632 | -2,28656 | 0,74731 |
| Srrm1 | -25,2604 | 2,80E-09 | -23,6789 | 1,16E-06 | -21,889 | 5,88E-06 |
| Srrm1 | -25,2604 | 2,80E-09 | -23,6789 | 1,16E-06 | -21,889 | 5,88E-06 |
| Sema6c | -25,7662 | 4,67E-07 | -7,25058 | 0,28041 | -23,2086 | 6,22E-05 |
| Sema6c | -25,7662 | 4,67E-07 | -7,25058 | 0,28041 | -23,2086 | 6,22E-05 |
| Mapk8ip3 | -26,0468 | 7,42E-35 | -3,17744 | 0,20138 | -2,58898 | 0,29512 |
| Gm8730 | -26,764 | 8,59E-21 | 16,93149 | 3,99E-07 | -1,45066 | 0,70601 |
| Thsd4 | -30 | 6,26E-09 | 2,429732 | 0,74018 | -0,03207 | 0,99637 |

**Supplementary table 3.** 100 most up and down regulated genes expressed in the injured sciatic nerve at P30, P4 and P10 vs uninjured sciatic nerve at P30, P4 and P10 respectively.

| | P30 | | P4 | | P10 | |
|---|---|---|---|---|---|---|
| **Pro-inflammatory macrophages** | | | | | | |
| **Gene ID** | **log2Fold Change** | **padj** | **log2Fold Change** | **padj** | **log2Fold Change** | **padj** |
| Arg1 | 5,501 | 3,3E-38 | 6,093 | 1,7E-36 | 5,378 | 6,0E-28 |
| Lgals3 | 4,704 | 2,6E-18 | 3,929 | 2,0E-10 | 3,231 | 1,8E-07 |
| Marco | 5,288 | 9,1E-08 | 5,065 | 5,3E-06 | 2,452 | 3,5E-02 |
| Ms4a4c | 5,089 | 5,6E-08 | 4,485 | 5,2E-05 | 5,912 | 2,1E-07 |
| Trem2 | 3,922 | 1,3E-08 | 3,351 | 1,7E-05 | 3,430 | 1,3E-05 |
| Ccl5 | 3,648 | 7,5E-08 | 1,910 | 1,4E-02 | 2,584 | 8,6E-04 |
| Ccr7 | 3,801 | 4,3E-06 | 2,686 | 3,4E-03 | 2,725 | 4,8E-03 |
| Cd38 | 1,348 | 3,9E-06 | 1,350 | 5,3E-05 | 1,307 | 8,2E-05 |
| Cd80 | 2,267 | 6,8E-05 | 1,711 | 1,7E-02 | 2,285 | 1,0E-03 |
| Cd86 | 0,309 | 5,8E-01 | 0,664 | 2,4E-01 | 0,338 | 5,7E-01 |
| Cxcl10 | 1,872 | 2,2E-04 | 1,314 | 2,7E-02 | 0,907 | 1,4E-01 |
| Fpr2 | 4,080 | 7,5E-04 | 3,863 | 5,0E-03 | 3,388 | 2,5E-02 |
| Il1b | 6,672 | 1,3E-10 | 6,711 | 1,8E-09 | 5,326 | 6,6E-05 |
| Il23a | -3,081 | 6,1E-13 | -2,815 | 2,2E-09 | -5,455 | 5,3E-11 |
| Il6 | 5,590 | 1,4E-01 | 5,412 | 2,1E-01 | 3,441 | 4,4E-01 |
| Inhba | 2,556 | 3,6E-07 | 2,145 | 6,8E-04 | 2,835 | 8,2E-07 |
| Nos2 | 1,884 | 1,1E-01 | 1,824 | 1,7E-01 | 3,448 | 6,0E-03 |
| S100a8 | 1,682 | 3,4E-01 | 4,807 | 6,1E-03 | 4,168 | 2,0E-02 |
| Socs1 | 2,182 | 2,0E-02 | 2,311 | 2,9E-02 | 2,223 | 3,6E-02 |
| Tlr2 | 2,110 | 3,1E-07 | 2,279 | 1,0E-06 | 1,884 | 6,6E-05 |
| Tlr4 | 1,258 | 5,6E-04 | 1,968 | 1,6E-06 | 1,344 | 1,1E-03 |
| Tnf | 3,965 | 1,1E-04 | 4,789 | 3,3E-05 | 2,325 | 5,7E-02 |
| **Anti-inflammatory macrophages** | | | | | | |
| Arg1 | 5,501 | 3,3E-38 | 6,093 | 1,7E-36 | 5,378 | 6,0E-28 |
| Cd163 | -1,087 | 1,7E-01 | -0,338 | 7,4E-01 | 0,222 | 8,3E-01 |
| Egr2 | -2,896 | 3,5E-15 | -3,901 | 3,7E-21 | -4,078 | 2,3E-23 |
| Fabp4 | 0,550 | 4,1E-01 | 1,343 | 4,3E-02 | 1,459 | 2,3E-02 |
| Fn1 | 1,985 | 5,0E-04 | 0,230 | 7,8E-01 | 3,487 | 1,9E-08 |
| Il10 | 3,674 | 5,7E-03 | 4,409 | 8,4E-04 | 2,868 | 8,6E-02 |
| Mfge8 | 1,499 | 8,0E-08 | 1,151 | 3,7E-04 | 1,178 | 2,3E-04 |
| Mrc1 | 0,355 | 6,4E-01 | 0,996 | 1,8E-01 | 1,414 | 4,4E-02 |
| Pdgfa | -0,321 | 5,7E-01 | -1,055 | 5,3E-02 | -0,279 | 6,5E-01 |
| Retnla | 2,386 | 4,9E-01 | 1,515 | 7,0E-01 | 2,187 | 5,5E-01 |
| Tgfb1 | 2,279 | 1,6E-21 | 1,570 | 1,1E-08 | 1,932 | 9,8E-13 |
| Vegfa | 0,481 | 6,1E-01 | 1,569 | 8,4E-02 | 1,291 | 1,6E-01 |
| **Repair Schwann cell** | | | | | | |
| Artn | 3,596 | 5,1E-05 | 3,795 | 1,0E-04 | 3,772 | 2,0E-04 |
| Ccl2 | 1,784 | 4,3E-04 | 3,932 | 1,1E-12 | 1,229 | 3,7E-02 |
| Shh | 6,279 | 6,2E-20 | 3,659 | 2,4E-05 | 1,668 | 4,7E-02 |
| Atf3 | 3,276 | 1,6E-28 | 2,709 | 9,5E-16 | 1,891 | 2,9E-08 |
| Cxcl12 | -0,362 | 3,2E-01 | -1,063 | 3,6E-03 | -0,214 | 6,1E-01 |
| Gap43 | 3,228 | 1,1E-15 | 2,374 | 2,6E-07 | 2,880 | 2,3E-10 |
| Gdnf | 7,635 | 3,4E-44 | 5,355 | 6,1E-17 | 5,698 | 4,0E-19 |

| | | | | | | |
|---|---|---|---|---|---|---|
| Gfap | -0,755 | 3,5E-01 | 1,381 | 1,2E-01 | -1,018 | 2,4E-01 |
| Id2 | -0,152 | 9,0E-01 | -0,494 | 6,7E-01 | -0,148 | 9,1E-01 |
| Junb | 2,775 | 8,2E-36 | 2,727 | 4,8E-27 | 2,697 | 1,4E-26 |
| Lif | 2,155 | 8,2E-02 | 1,522 | 3,0E-01 | 0,993 | 5,1E-01 |
| Ncam1 | 1,621 | 5,3E-02 | 1,262 | 1,9E-01 | 2,843 | 1,2E-03 |
| Ngf | 0,650 | 6,3E-01 | 1,637 | 2,2E-01 | 0,619 | 6,7E-01 |
| Ngfr | 2,878 | 2,7E-14 | 0,471 | 3,5E-01 | 0,804 | 8,2E-02 |
| Olig1 | 7,943 | 3,5E-14 | 6,113 | 1,2E-27 | 7,280 | 2,4E-13 |
| Osm | 2,072 | 3,3E-03 | 3,592 | 2,9E-06 | 2,031 | 2,0E-02 |
| S100b | -2,402 | 2,9E-08 | -3,788 | 3,5E-15 | -4,340 | 6,7E-20 |
| Sox2 | 2,181 | 6,3E-10 | -0,083 | 8,8E-01 | 0,164 | 7,5E-01 |

**Supplementary table 4**. Representative Pro-inflammatory anti-inflammatory macrophage and repair Schwann cell genes expressed in the injured sciatic nerve at P30, P4 and P10 vs uninjured sciatic nerve at P30, P4 and P10 respectively.
